# A reversible GNAS-PKA-RUNX2 checkpoint governs skeletal stem cell fate in fibrous dysplasia

**DOI:** 10.64898/2026.10.02.756293

**Authors:** Elena Sofia Cardenas-Alcoser, Diego Rodriguez, Angelyn Solis, Kuniaki Sato, Tomohiko Ishikawa, Sendi Rafael Adame-Garcia, Tsubasa Ochiai, Johannes Koeckenberger, Soo Hyun Shin, Jiang Du, Fleur Ferguson, J Silvio Gutkind

## Abstract

Fibrous dysplasia (FD) arises from activating *GNAS* mutations that increase cAMP signaling in skeletal stem/progenitor cells (SSPCs), but the underlying mechanisms and whether the resulting fate imbalance can be reversed remain unclear. Using controlled *GNAS* induction with transcriptomic, pharmacologic, and cell-fate analyses, we identify a PKA-dependent checkpoint governing SSPC lineage commitment. *GNAS*^R201C^ redirects SSPCs toward osteoclastogenesis while suppressing osteoblast maturation; genetic or pharmacologic PKA inhibition reverses both effects, revealing this fate decision is dynamically reversible. Mechanistically, PKA phosphorylates RUNX2, the master regulator of osteogenesis, enforcing its cytoplasmic retention and blocking osteogenic transcription; a non-phosphorylatable RUNX2 mutant restores nuclear localization and osteoblast differentiation, defining RUNX2 phosphorylation as a disease-initiating molecular switch. Critically, PKA inhibition in vivo reduces fibrosis and osteoclast burden in established bone lesions, demonstrating reversibility even in advanced FD. These findings define a druggable GNAS-PKA-RUNX2 checkpoint and identify PKA inhibition as a strategy to reverse existing disease in FD.

**Graphical abstract:** 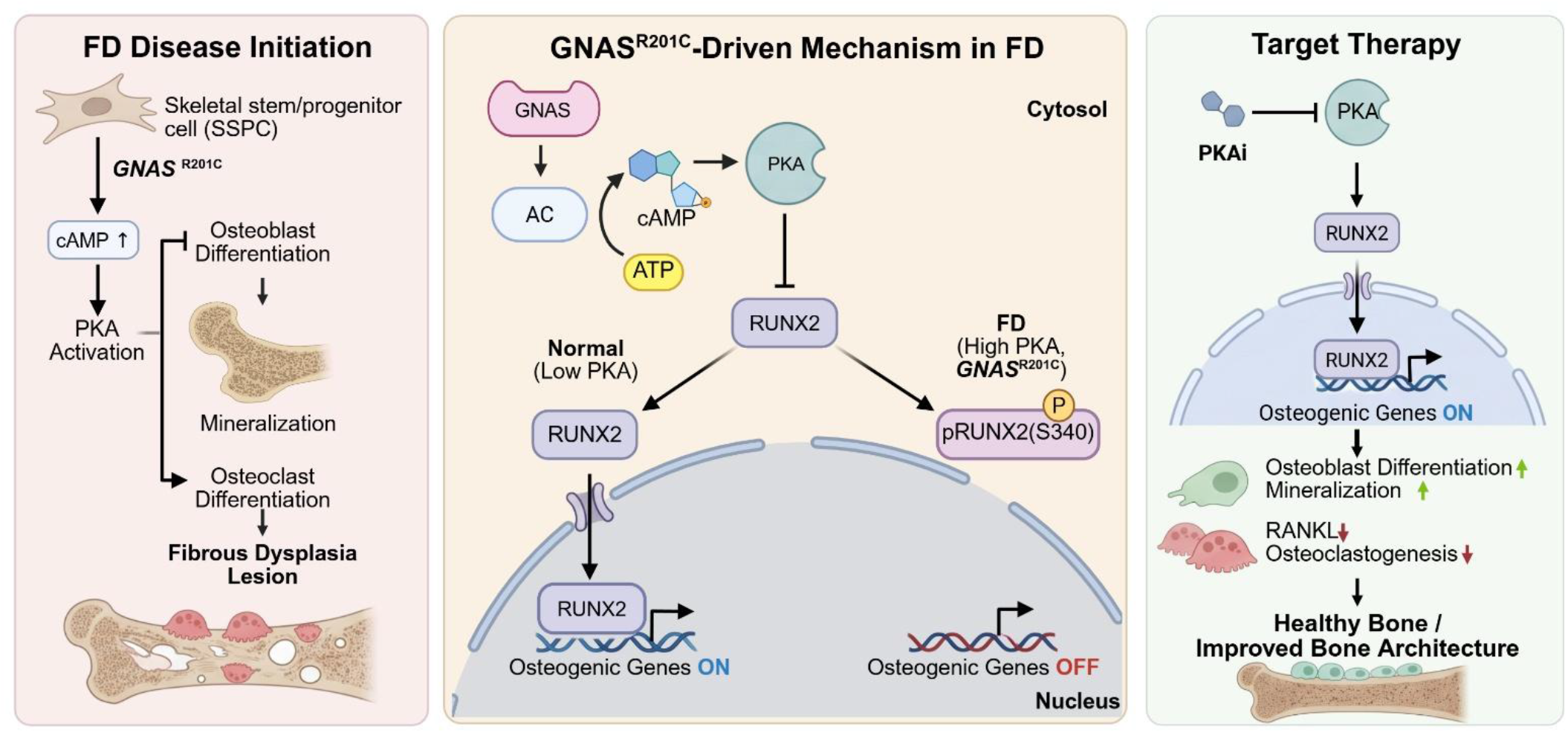

## Background

Fibrous dysplasia (FD) is a mosaic skeletal disorder in which normal lamellar bone is replaced by fibro-osseous tissue, resulting in bone deformity, pain, and pathologic fractures^1^. FD is driven by post-zygotic mutations in *GNAS*, most commonly R201C/H, that encode a constitutively active Gαs, elevating cAMP and chronically stimulating the protein kinase A (PKA) holoenzyme^2^. In skeletal stem and progenitor cells (SSPCs), persistent cAMP-PKA signaling disrupts osteolineage differentiation and arrests osteoblast maturation. It establishes an aberrant pro-osteoclastogenic environment, thereby disrupting the balance between bone formation and resorption^3, 4^. Despite a well-defined genetic lesion and pathway dependency, no approved precision therapies exist for FD; current management remains largely supportive and anti-resorptive, such as denosumab, which blocks RANKL activity, rather than disease-modifying^5, 6^.

A principal obstacle to therapeutic progress has been the absence of tractable, temporally controlled models that recapitulate FD pathophysiology from SSPCs through tissue remodeling and disease progression. To address this problem, we engineered a tetracycline-inducible *GNAS*^R201C^ mouse model (FD model) that faithfully recapitulates the histologic hallmarks of FD within two weeks of transgene induction^4^. Similarly, constitutive activation of PKA-Cα (*PRKACA*) subunit was sufficient to generate FD-like lesions, while genetic or pharmacological PKA suppression significantly reduced osteoclastogenesis and prevented the initiation of FD^7^. Additionally, our prior studies showed that doxycycline withdrawal triggers regression of established lesions and restoration of bone architecture, indicating that sustained *GNAS*–PKA hyperactivation is required for lesion maintenance in addition to disease initiation^4^. Collectively, these findings suggest that the hyperactivation of the *GNAS*-PKA signaling axis is a key driver of FD pathogenesis. However, the downstream mechanisms linking PKA activity to osteogenic dysregulation remain poorly defined. In particular, how PKA signaling interfaces with key drivers of osteolineage differentiation has not been fully elucidated. Specifically, the molecular events linking PKA malfunction to failure in osteoblast maturation and excess osteoclastogenic signaling in SSPCs remain incompletely understood^3^. Identifying the components that mediate this connection would reveal novel druggable nodes for disease modification rather than symptomatic palliation.

The recently described small molecule BLU0588 is a highly selective PRKACA inhibitor that effectively suppresses PKA signaling in fibrolamellar carcinoma models^8^. In vivo, oral administration of BLU0588 produces rapid, sustained pathway inhibition, suppressing PKA-driven tumor growth without significant toxicity, as evidenced by stable body weight^8^. Complementing these preclinical findings, our team’s recent structural characterization of the *PRKACA*-BLU0588 complex revealed the molecular basis for the compound’s selectivity, demonstrating stabilization of a distinct open yet ordered kinase conformation and further supporting the therapeutic targeting of PKA-driven diseases^9^. Together, these studies further support *PRKACA* as a therapeutically actionable target and highlight the potential of BLU0588 to pharmacologically inhibit a kinase long considered “undruggable.”

Here, we integrate the study of inducible *in vitro* and *in vivo* FD models, pathway-targeted chemical biology, transcriptomics, and mechanistic analysis to identify the central molecular components through which hyperactive *PRKACA* drives FD development. We identify RUNX2, a master regulator of osteoblast differentiation, as a critical substrate of PKA signaling and demonstrate that its phosphorylation rewires osteogenic programs to promote FD pathology. Furthermore, we show that pharmacological inhibition of PKA activity restores established lesions, highlighting the reversibility of disease-associated cellular states. Together, these findings establish a direct mechanistic link between aberrant PKA signaling and skeletal stem cell osteogenic dysregulation and provide a framework for targeted therapeutic intervention to halt key FD signaling drivers.

## Results

### 1. Establishment of an inducible bone marrow stem cell (BMSC)-Derived osteolineage model to assess *GNAS*^R201C^ function

To interrogate how *GNAS*-driven signaling alters osteolineage differentiation, we established a controlled in-vitro system using bone marrow–derived stromal cells (BMSCs) isolated from our tetracycline-inducible *Prrx1*-CRE/LSL-rTtA-EGFP/TetON-*GNAS*^R201C^/TetON-H2B-GFP (FD) mouse model. After isolation and 10-day expansion, BMSCs were exposed for 48 hours to doxycycline (Dox) to induce Gαs^R201C^ expression (**Fig. 1A**). A dose–response curve identified 2 μM Dox as the most effective and reproducible condition to induce Gαs protein expression (**Fig. 1B**). Nuclear expression of the H2B-GFP fluorescent reporter validated the efficiency of the Tet-On system (**Fig. 1C**).

**Figure 1.**
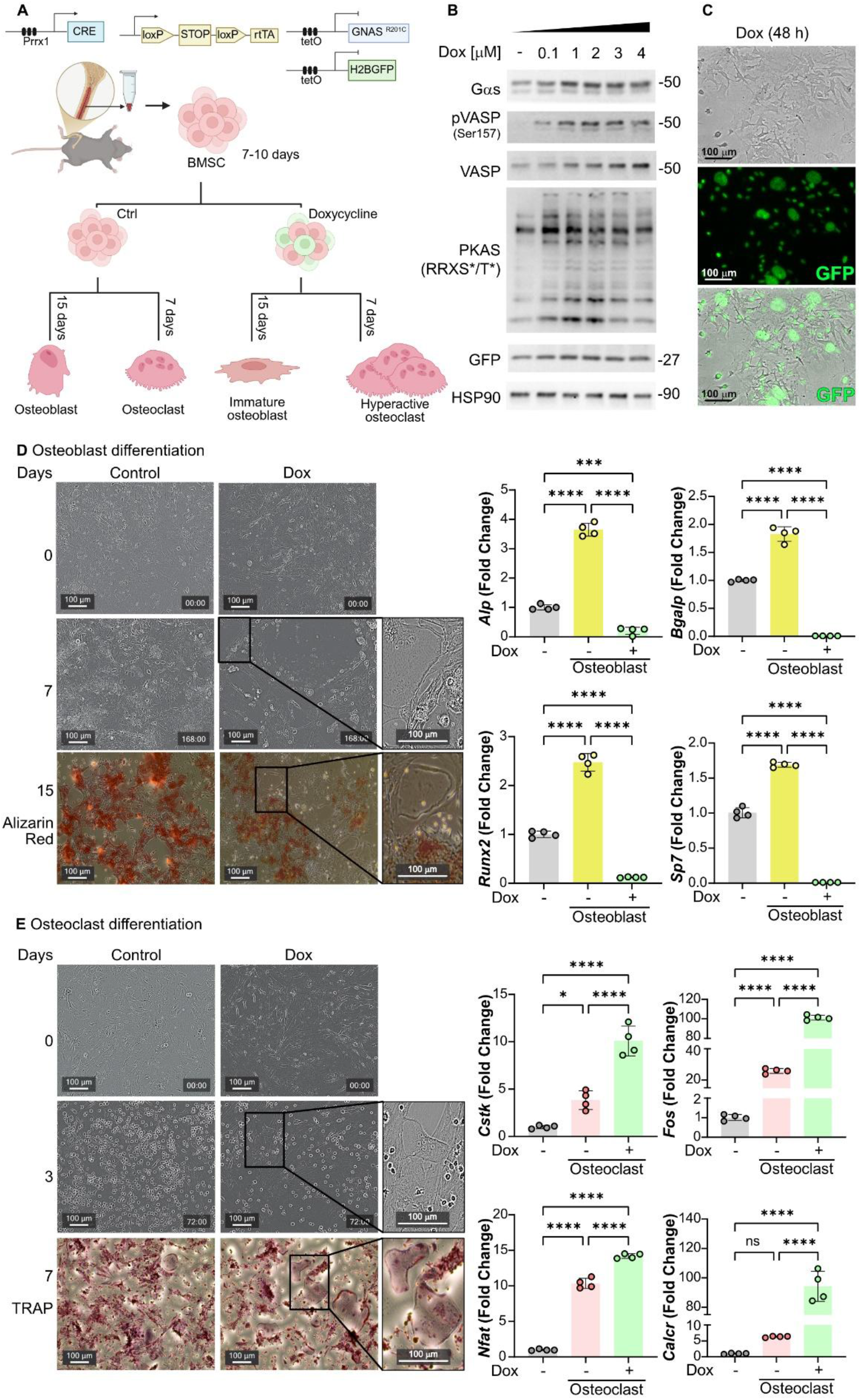
Inducible GNAS^R201C^ activation in BMSC suppresses osteoblast differentiation and enhances osteoclastogenesis. **A**. Schematic of the experimental strategy. BMSCs were isolated from *Prrx1*-Cre;LSL-rtTA;tetO-GNAS^R201C^;tetO-H2BGFP (FD model) mice and expanded in culture for 7–10 days before doxycycline (Dox)-induced transgene activation. Cells were subsequently differentiated toward the osteoblast or osteoclast lineage in the absence or presence of Dox. Osteoblast differentiation was assessed after 15 days, whereas osteoclast differentiation was evaluated after 7 days. Created in BioRender. **B**. Immunoblot analysis of BMSCs treated for 48 h with increasing concentrations of Dox (0.1–4 μM). Expression of Gαs, phosphorylation of the PKA target VASP (Ser157), pan-phospho-PKA substrates (RRXS/T), and GFP are shown. HSP90 served as a loading control. **C**. Representative brightfield and fluorescence images of BMSCs following 48 h of Dox treatment. Scale bars, 100 μm. **D**. Osteoblast differentiation of control and Dox-treated BMSCs. Representative phase-contrast images at days 0 and 7 and Alizarin Red staining at day 15. Higher-magnification views highlight cellular morphology. Quantitative RT-PCR analysis of osteoblast-associated genes (*Alp, Bgalp, Runx2*, and *SP7*) in control and Dox-treated cultures. Data are presented as fold change relative to control cultures. **E**. Osteoclast differentiation of control and Dox-treated BMSCs. Representative phase-contrast images at days 0 and 3 and TRAP staining at day 7. Enlarged views show multinucleated osteoclast-like cells. Quantitative RT-PCR analysis of osteoclast-associated genes (*Ctsk, Fos, Nfatc1*, and *Calcr*) in control and Dox-treated cultures. Data are presented as fold change relative to control cultures. Data are shown as mean ± SD. Each dot represents an independent biological replicate. Statistical significance was determined using one-way ANOVA with multiple-comparison correction. *P < 0.05, ***P < 0.001, ****P < 0.0001; ns, not significant.

Under osteogenic conditions (i.e., β-glycerol phosphate, ascorbic acid, and a dexamethasone-rich medium^10^), control cultures progressed to mature mineralization by day 15, as assessed by increased alizarin red signal. In contrast, *GNAS*^R201C^ activation redirected the SSPCs’ differentiation trajectory: multinucleated osteoclast-like cells emerged by day 7 and became prominent by day 15, accompanied by reduced staining of calcium deposition. qPCR analysis demonstrated suppression of key osteoblast differentiation and maturation factors (e.g., *Runx2*, *Sp7, Alp*, and *Bglap*) (**Fig. 1D**). In parallel, osteoclast differentiation conditions (e.g., using RANKL and m-CSF-rich medium^11^) showed accelerated osteoclastogenesis upon *GNAS*^R201C^ expression, with functional osteoclasts appearing as early as day 3. In control conditions, on day 3 only pre-osteoclasts are present, and multinuclear cells develop later, on day 7. *GNAS*^R201C^ expression also increased the expression of mature osteoclast-associated genes (e.g., *Ctsk, Calcr, Nfat, Fos*) (**Fig. 1E**). Together, these findings establish a tractable inducible platform to model *GNAS*^R201C^ effects in osteolineage cells and demonstrate that mutant Gαs expression is sufficient to initiate changes in the BMSC fate, halting osteoblast maturation and promoting exaggerated osteoclastogenesis.

### 2. PKA catalytic subunit α (PKA-Cα) knockdown restores osteolineage gene expression in *GNAS*^R201C^ BMSCs

PKA is organized as a heterotetrameric holoenzyme assembled by two regulatory (RIα, RIβ, RIIα, and RIIβ) and two catalytic (Cα, Cβ, and Cγ) subunits, which together determine the spatial and functional specificity of PKA signaling^12^. Among the catalytic subunits, PKA-Cα (encoded by *PRKACA*) is the principal catalytic isoform implicated in osteoblast function and bone homeostasis, mediating many of the downstream effects of cAMP-PKA signaling in skeletal tissues^13, 14^. To determine whether the osteolineage defects induced by *GNAS*^R201C^ are driven specifically by downstream overactivation of the PKA-Cα, we performed a knockdown of *Prkaca* in BMSCs using siRNAs. BMSCs from our inducible *GNAS*^R201C^ mouse model were transfected with control or PKA-Cα siRNAs, followed by Dox-induced mutant Gαs expression (**Fig. 2A-B**). First, we confirmed that knockdown of PKA-Cα did not alter the expression levels of the inducible *GNAS* mutant (**Fig. 2C**), indicating that the siRNA intervention selectively modulated PKA signaling without interfering with Tet-On–driven transgene induction. We next evaluated how suppression of PKA activity affected the osteolineage transcriptional program under GNAS^R201C^ activation. Dox-induced cells transfected with the control siRNA exhibited increased expression of osteoclastogenic regulators in immature osteoblasts, consistent with the aberrant pro-osteoclast environment observed in FD, as shown in Figure 1. However, knockdown of PKA-Cα (KD PKA-Cα) significantly reduced the expression of RANKL and increased the ratio RANKL/OPG (**Fig. 2D**), indicating that excessive PKA activity is required for their induction. Concomitantly, suppression of PKA-Cα produced a similar effect on osteoblast differentiation markers. In *GNAS*^R201C^-expressing cells, KD PKA-Cα resulted in a subtle rescue of osteoblast-associated genes (**Fig. 2E**), while osteoclast lineage markers were markedly decreased (**Fig. 2F**).

**Figure 2.**
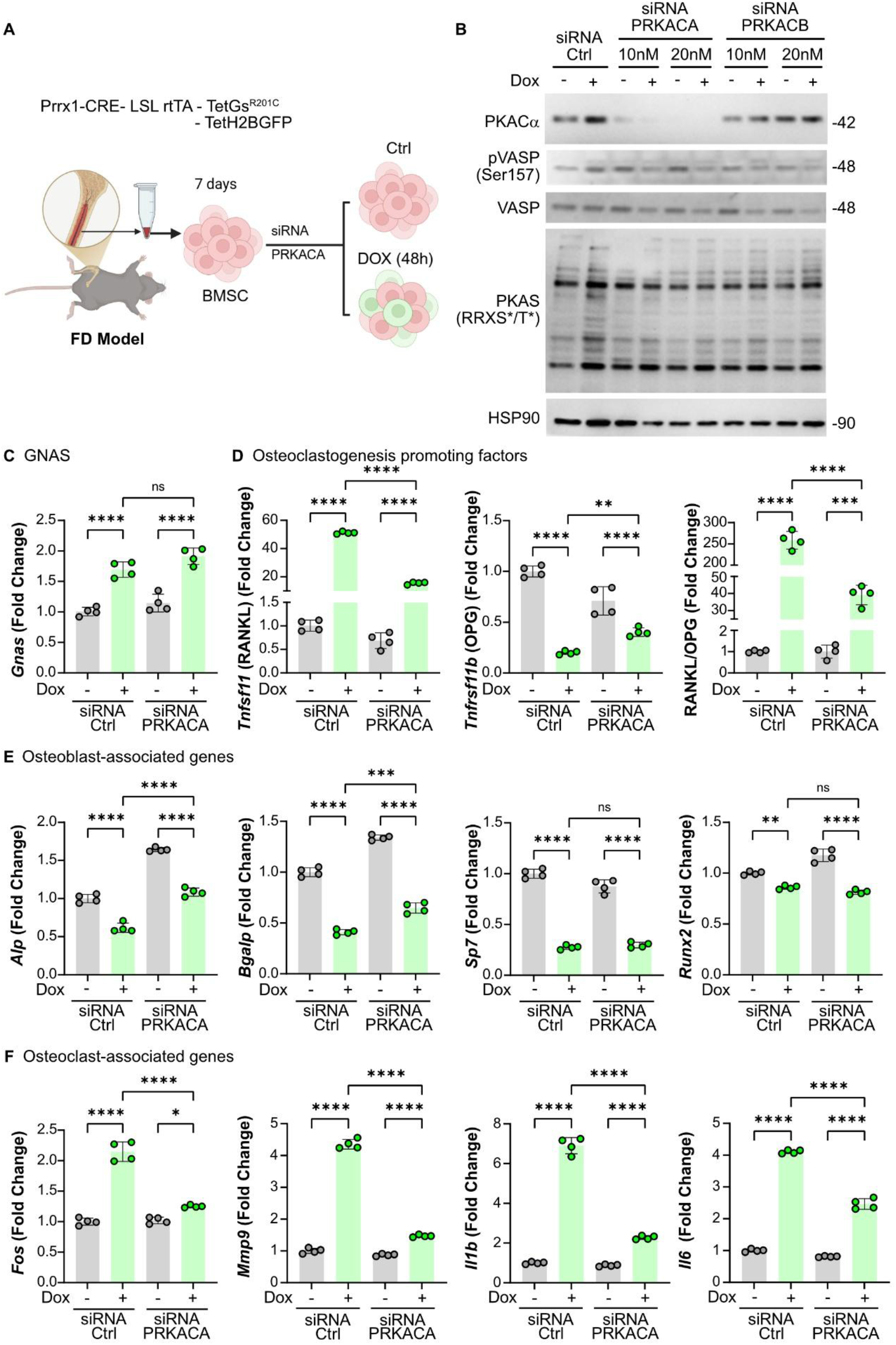
PKAα is a critical downstream effector of GNAS^R201C^ that drives osteoclastogenic signaling and represses osteoblast gene expression programs. **A**. Experimental design. BMSCs isolated from FD mice were cultured for 7 days and transfected with control (Ctrl) or *Prkaca* siRNA prior to induction of GNAS^R201C^ expression with doxycycline (Dox) for 48 h. Created in BioRender. **B.** Immunoblot analysis of BMSCs transfected with control siRNA or increasing concentrations of siRNAs targeting PRKACA or PRKACB and treated with Dox for 48 h. Protein levels of PKA-Cα, phosphorylated VASP (Ser157), total VASP, and phosphorylated PKA substrates (RRXS/T) were assessed. HSP90 served as a loading control. **C.** Quantitative RT-PCR analysis of C. *Gnas,* D. osteoclastogenesis-promoting factors (*Tnfsf11* (RANKL), *Tnfrsf11b* (OPG), and the RANKL/OPG), **E.** osteoblast-associated genes (*Alp, Bgalp, Sp7*, and *Runx2*), **F.** osteoclast-associated and pro-inflammatory genes (*Fos, Mmp9, Il1b,* and *Il6*) ratio expression in control and *Prkaca*-silenced BMSCs following Dox treatment. Data are presented as mean ± SD. Each dot represents an independent biological replicate. Statistical significance was determined by one-way ANOVA with multiple-comparison correction. ns, not significant; *P < 0.05, **P < 0.01, ***P < 0.001, ****P < 0.0001.

Together, these changes demonstrate that inhibiting PKA catalytic activity can restore the balance of osteolineage gene expression disrupted by mutant *GNAS*. Reduced PKA-Cα expression decreased osteoclastogenic signaling while promoting recovery of osteoblast differentiation markers, further supporting a role for PKA signaling in mediating the osteolineage defects associated with mutant *GNAS*.

### 3. Pharmacologic PKA inhibition restores osteolineage transcriptional programs in *GNAS*^R201C^ BMSCs

To further evaluate whether PKA catalytic activity is required for the osteolineage abnormalities induced by *GNAS*^R201C^, we treated FD-BMSCs with the selective small-molecule PKA inhibitor BLU0588 (PKAi), a potent *PRKACA* inhibitor that allows direct pharmacological targeting of catalytic PKA activity^8, 9^. Western blot analyses demonstrated that PKAi effectively suppressed PKA pathway activity, as evidenced by a marked reduction in VASP phosphorylation, a typical PKA target (**Fig. 3A**), and a global decrease in PKA-phosphorylated substrates, as assessed using a phospho-PKA substrate antibody (PKAS), which recognizes serine/threonine residues phosphorylated within the consensus PKA motif (RRXS*/T*). This antibody detects a broad range of endogenous PKA targets, providing a readout of overall PKA-dependent phosphorylation. Accordingly, changes in signal intensity reflect global alterations in canonical PKA substrate phosphorylation. PKA activity was also tested using a cAMP-response element (CRE)-luciferase reporter assay to further validate on-target transcriptional inhibition: PKAi treatment significantly reduced CRE-driven transcriptional activity upon Dox induction (**Fig. 3C**), demonstrating effective blockade of cAMP-PKA signaling at the transcriptional level. Importantly, inhibition of PKA did not alter Dox-induced expression of the *GNAS* mutant at either the protein or transcript level (**Fig. 3A-B**), confirming that PKAi targets the downstream PKA arm of the pathway without interfering with the inducible transgene. As a control, the same response was observed with Gs-coupled GPCR reporter activation, using HEK293A cells with a safe harbor locus (AAVS1)-targeted Gs-Designer Receptors Exclusively Activated by Designer Drugs (DREADD) knock-in. We found that PKAi blocks CNO-induced activation of the PKA pathway and significantly decreases CRE-driven reporter signaling (**Supplementary Fig 1A-B**), supporting a general signaling effect of BLU0588 rather than exhibiting cell-specific activity.

**Figure 3.**
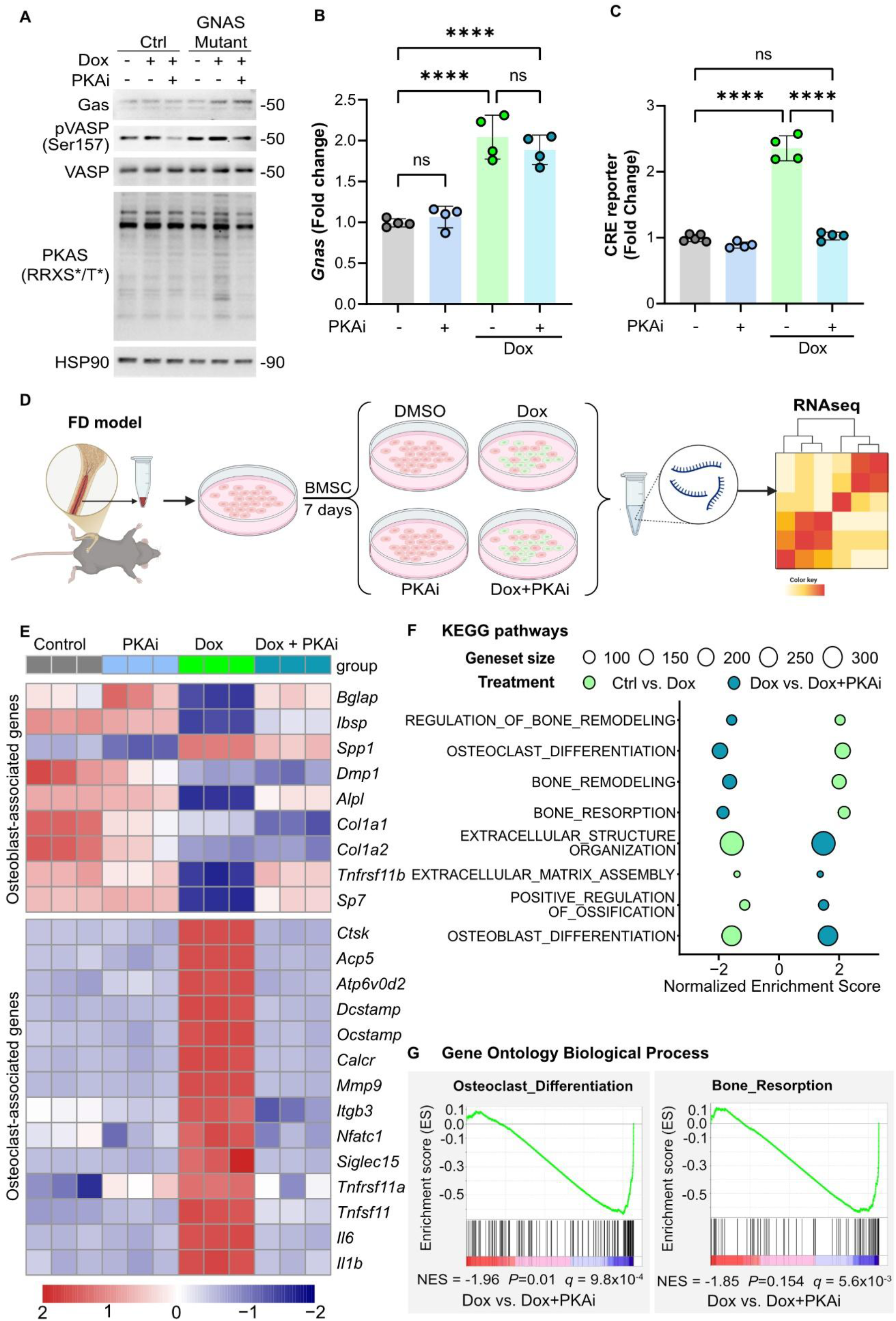
Pharmacological inhibition of PKA reverses GNAS^R201C^-driven transcriptional programs in FD BMSCs. **A.** Immunoblot analysis of control and GNAS^R201C^-expressing BMSCs treated with doxycycline (Dox) in the presence or absence of a PKA inhibitor (PKAi). Protein levels of Gαs, phosphorylated VASP (Ser157), total VASP, and phosphorylated PKA substrates (RRXS/T) were assessed. HSP90 served as a loading control. **B.** Quantitative RT-PCR analysis of *Gnas* expression in control and GNAS^R201C^ BMSCs treated with PKAi and Dox. **C.** CRE-luciferase reporter activity in GNAS^R201C^ BMSCs treated with Dox in the presence or absence of PKAi. Data are presented as fold change relative to untreated controls. Renilla expression was used as internal normalization control. **D.** Experimental design. BMSCs isolated from FD mice were cultured for 7 days and treated with vehicle (DMSO), Dox, PKAi, or Dox+PKAi before RNA isolation and transcriptomic analysis by RNA sequencing. Created in BioRender. **E.** Heatmap showing expression of osteoblast-associated genes and osteoclast-associated genes across treatment conditions. **F.** Gene set enrichment analysis (GSEA) comparing Dox- and Dox+PKAi-treated FD BMSCs. Bubble plot showing significantly enriched biological pathways. **G.** Representative enrichment plots for the osteoclast differentiation and bone resorption gene sets are shown. Data are presented as mean ± SD. Each dot represents an independent biological replicate. Statistical significance was determined by one-way ANOVA with multiple-comparison correction. ns, not significant; ****P < 0.0001.

To determine how PKA inhibition reshapes the osteolineage transcriptional program, we performed bulk RNA sequencing (**Fig. 3D**). *GNAS*^R201C^ activation resulted in a pronounced upregulation of osteoclast-associated gene programs, consistent with the aberrant pro-resorptive signature observed in FD (**Fig. 3E**). Of note, treatment with PKAi markedly reversed this aberrant transcriptional pattern, strongly diminishing osteoclastogenic gene expression. Simultaneously, PKA inhibition rescued osteoblast gene expression, supporting a shift toward an osteogenic transcriptional identity (**Fig. 3E**). Consistent with these observations, gene set enrichment analysis (GSEA) indicated the negative enrichment of bone development and ossification pathways in the presence of *GNAS*^R201C^, which were significantly reactivated upon PKAi treatment, suggesting that PKA inhibition may reprogram mutant-*GNAS*-induced transcription. (**Fig. 3F and 3G**).

To functionally validate the transcriptional changes identified in our RNA-seq analysis, we next examined whether PKA inhibition could restore osteoblast maturation and attenuate osteoclastogenic signals at the cellular and biochemical levels. Consistent with the transcriptomic rescue observed with PKAi, qPCR analysis confirmed that typical osteoblast differentiation genes (e.g., *Bgalp*, *Sp7, Tnfrs11b*, and *Runx2*) were partially restored upon PKA inhibition in *GNAS*^R201C^-expressing BMSCs (**Fig. 4A**). In addition, osteoclastogenic regulators revealed the opposite trend: key pro-osteoclast factors were significantly reduced following PKA inhibition (**Fig. 4B-C**), indicating a shift away from the pathological osteoclastogenic environment. To determine whether this transcriptional shift translated into altered paracrine signaling, we measured the release of RANKL into the conditioned media. PKAi treatment markedly decreased secreted RANKL (**Fig. 4D**), consistent with the reduction in osteoclastogenesis-promoting transcripts and suggesting that PKA activity directly influences osteoblast-derived signals that control osteoclast differentiation. Functionally, these molecular changes produced a measurable effect on osteoclast behavior. Expression of *GNAS*^R201C^ in BMSCs robustly stimulated osteoclast differentiation and activity in a fluorescent calcium-phosphate bone resorption assay, whereas PKAi-treated cultures induced significantly lower resorption activity (**Fig. 4F**), demonstrating that PKA inhibition reduces the pro-osteoclastogenic capacity of *GNAS*^R201C^ BMSCs. Finally, analysis of osteoclast-specific genetic markers (e.g., *Cstk, Mmp9, Il6*, and *Il1b*) showed that all major osteoclast markers returned toward baseline levels after PKAi treatment (**Fig. 4G**), confirming that pharmacologic PKA inhibition reverses the osteoclastogenic signaling cascade initiated by mutant *GNAS*. Consistent with the transcriptional alterations induced by aberrant PKA signaling, these were mirrored at the protein level, as expression of osteoblast and osteoclast markers was normalized following PKA inhibition (**Supplementary Fig 1C**).

**Figure 4.**
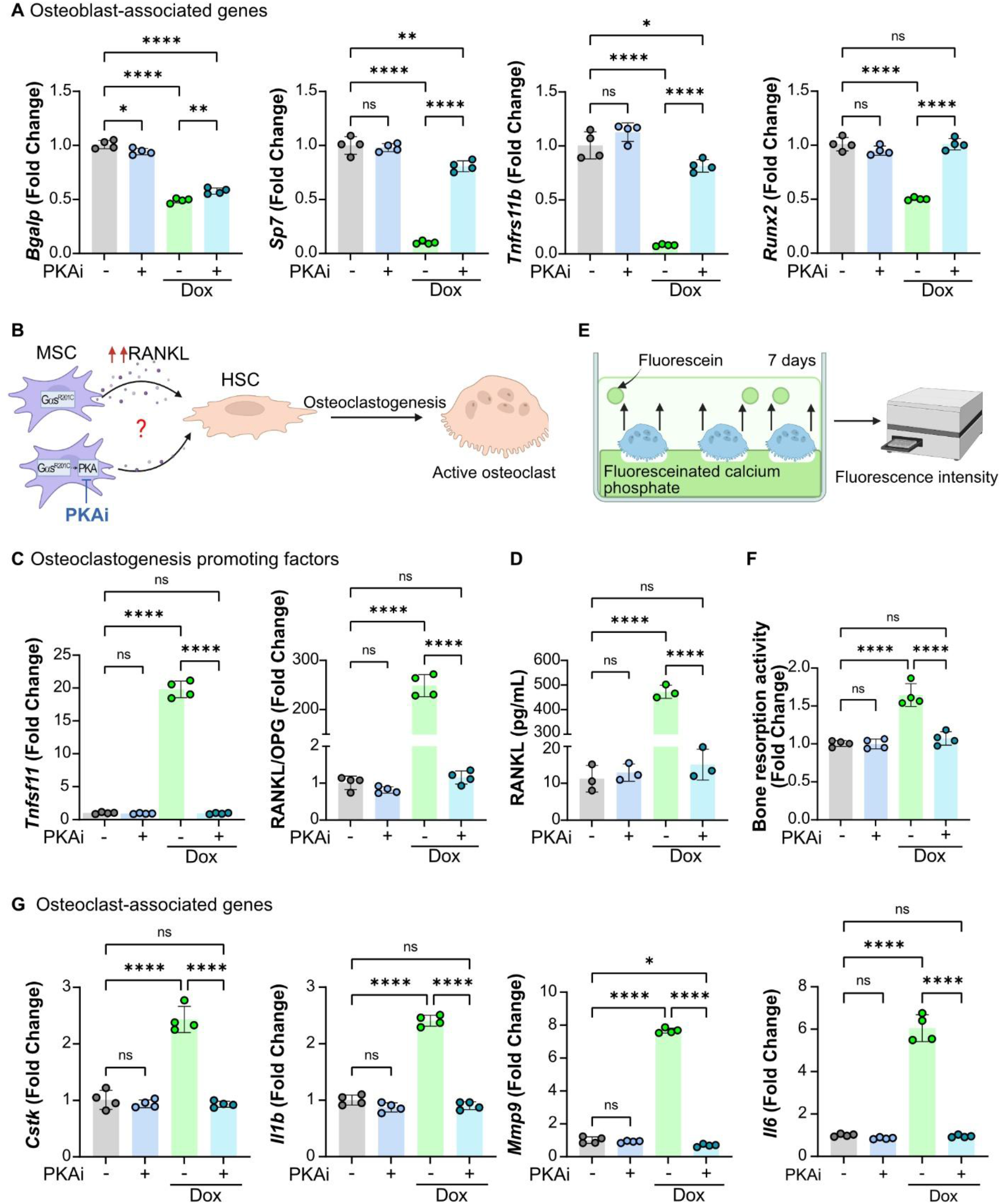
PKA inhibition suppresses FD BMSC-derived osteoclastogenic signaling and osteoclast activation. **A.** Quantitative RT-PCR analysis of osteoblast-associated genes (*Bglap, Sp7, Tnfrsf11b,* and *Runx2*) in FD BMSCs treated with doxycycline (Dox) in the presence or absence of a PKA inhibitor (PKAi). **B.** Proposed model for indirect regulation of osteoclastogenesis by GNAS^R201C^-expressing mesenchymal stromal cells (MSCs). Created in BioRender. **C.** Quantitative RT-PCR analysis of osteoclastogenesis-promoting factors, including *Tnfsf11* (RANKL) and the RANKL/OPG ratio, in FD BMSCs treated with Dox in the presence or absence of PKAi. **D.** Quantification of secreted RANKL protein levels in conditioned media from FD BMSCs treated with Dox and/or PKAi, as determined by ELISA. **E.** Schematic of the in vitro bone resorption assay. Created in BioRender. **F.** Quantification of osteoclast-mediated bone resorption activity of FD BMSCs treated with Dox and/or PKAi. **G.** Quantitative RT-PCR analysis of osteoclast-associated and inflammatory genes (*Ctsk, Il1b, Mmp9,* and *Il6*) in osteoclast cultures exposed to conditioned media from FD BMSCs treated with Dox and/or PKAi. Data are presented as mean ± SD. Each dot represents an independent biological replicate. Statistical significance was determined by one-way ANOVA with multiple-comparison correction. ns, not significant; *P < 0.05, **P < 0.01, ****P < 0.0001.

These functional assays validate our transcriptomic findings and demonstrate that PKA inhibition not only rebalances osteolineage gene expression but also normalizes osteoblast output, reduces RANKL-mediated osteoclastogenesis, and suppresses osteoclast functional activity. These results establish that PKAi effectively improves both the molecular and functional consequences of *GNAS*^R201C^-PKA-driven osteolineage dysregulation.

### 4. PKA inhibition normalizes the transcriptional and functional profile of mature osteoclasts differentiated from *GNAS*^R201C^ BMSCs

Building on the differentiation framework established above, we next asked whether pharmacologic PKA inhibition could normalize the osteoclast overactivity observed under *GNAS*^R201C^ expression. First, BMSCs from our inducible model were differentiated into mature osteoclasts under Dox, with or without PKAi treatment (**Fig. 5A**). Bulk RNA-seq of mature osteoclast cultures confirms that *GNAS*^R201C^ results in overactivation of osteoclast-associated genes. Treatment with PKAi reversed this transcriptional pattern, restoring expression of canonical osteoclast markers toward baseline (**Fig. 5B**). Pathway-level analyses were concordant: signatures linked to bone remodeling, osteoclast differentiation, bone resorption, extracellular structure organization/ECM assembly, and positive regulation of ossification, as well as osteoblast differentiation pathways, were all returned toward control levels when PKA activity was inhibited (**Fig. 5C**).

**Figure 5.**
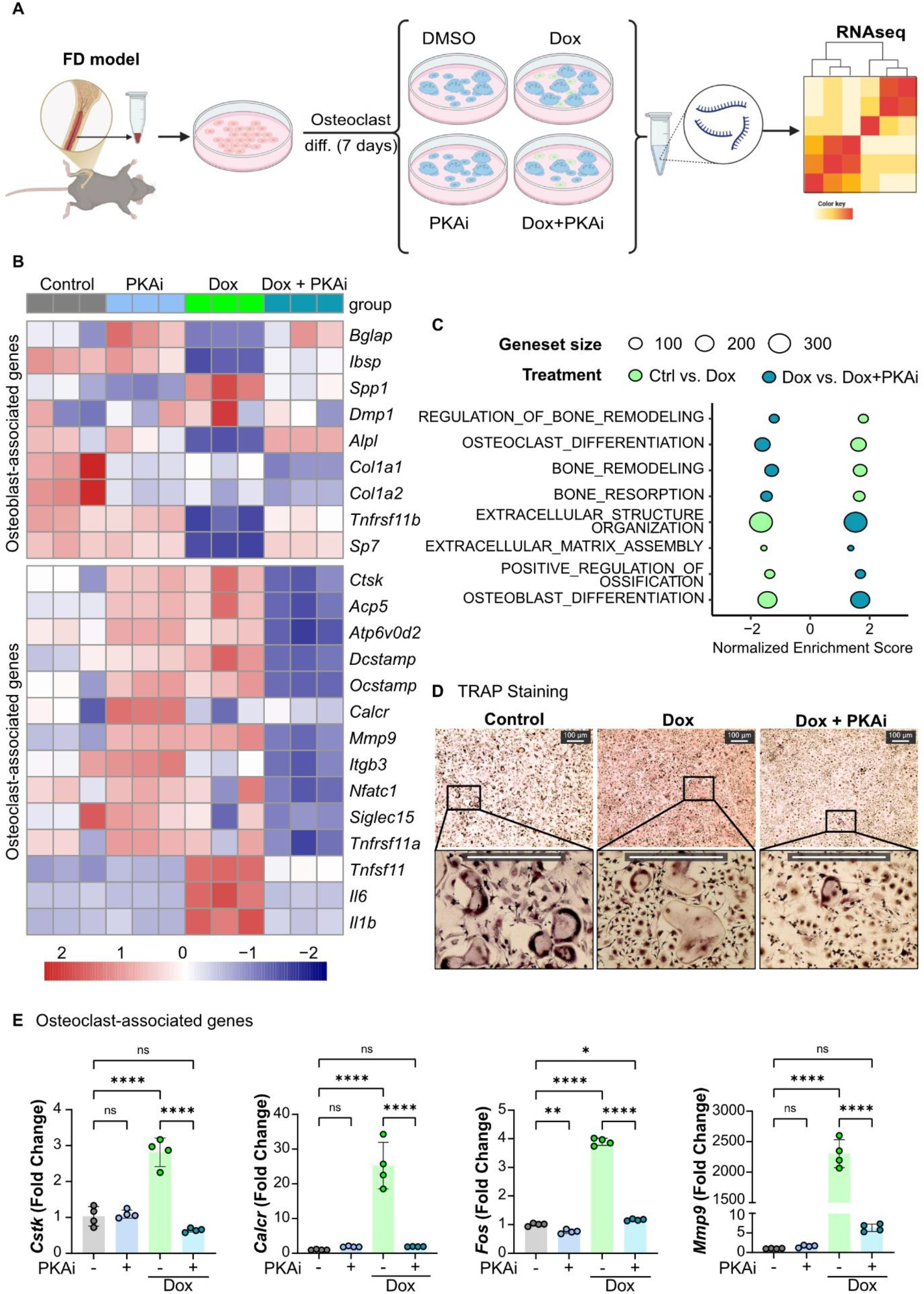
PKA inhibition reverses the osteoclastogenic transcriptional program induced by GNAS^R201C^ during osteoclast differentiation. **A.** Experimental design. BMSCs isolated from FD mice were differentiated toward the osteoclast lineage for 7 days in the presence of vehicle (DMSO), doxycycline (Dox), PKA inhibitor (PKAi), or Dox+PKAi prior to RNA isolation and transcriptomic analysis by RNA sequencing. Created in BioRender. **B.** Heatmap showing expression of osteoblast-associated genes and osteoclast-associated genes across treatment conditions. **C.** Gene set enrichment analysis (GSEA) comparing Dox- and Dox+PKAi-treated osteoclast cultures. Bubble plot showing significantly enriched biological pathways. **D.** Representative TRAP staining of osteoclast cultures treated with vehicle, Dox, or Dox+PKAi. Insets show higher magnification views of multinucleated TRAP-positive osteoclasts. Scale bars, 100 μm. **E.** Quantitative RT-PCR analysis of osteoclast-associated genes (*Ctsk, Calcr, Fos,* and *Mmp9*) in osteoclast cultures treated with Dox in the presence or absence of PKAi. Data are presented as mean ± SD. Each dot represents an independent biological replicate. One-way ANOVA determined statistical significance with multiple-comparison correction. ns, not significant; *P < 0.05, **P < 0.01, ****P < 0.0001.

We next assessed whether transcriptional normalization is translated into functional rescue. Cellular analyses supported these findings: TRAP staining showed fewer TRAP-positive multinucleated cells in PKAi-treated cultures compared with Dox alone (**Fig. 5D**), indicating a reduced osteoclast burden under PKA blockade. Finally, qPCR confirmed the decrease of osteoclast-specific genes (e.g., *Ctsk, Calcr, Fos,* and *Mmp9*) toward baseline after PKAi treatment (**Fig. 5E**), aligning with the RNA-seq and functional readouts.

These findings demonstrate that PKA inhibition normalizes both the transcriptional identity and functional behavior of mature osteoclasts derived from *GNAS*^R201C^ BMSCs, reinforcing that excessive PKA catalytic signaling represents the proximal driver of *GNAS*-induced osteoclast overactivation in this model.

### 5. PKA inhibition restores osteoblast differentiation and suppresses osteoclastogenic signaling in *GNAS*^R201C^ BMSCs

To determine whether pharmacologic inhibition of PKA can directly rescue the osteoblast differentiation defects caused by *GNAS*^R201C^, we used the same model as above, with or without PKAi (**Fig. 6A**). Bulk RNA-seq revealed that Dox induction resulted in a marked upregulation of osteoclast-associated genes within the osteoblast-differentiation-directed cultures, consistent with the establishment of an aberrant pro-osteoclastogenic state. Treatment with PKAi abolished this induction, returning the osteoclast markers to basal levels. In parallel, osteoblast-associated markers exhibited a partial restoration with PKA inhibition, supporting reactivation of the osteogenic transcriptional program (**Fig. 6B**).

**Figure 6.**
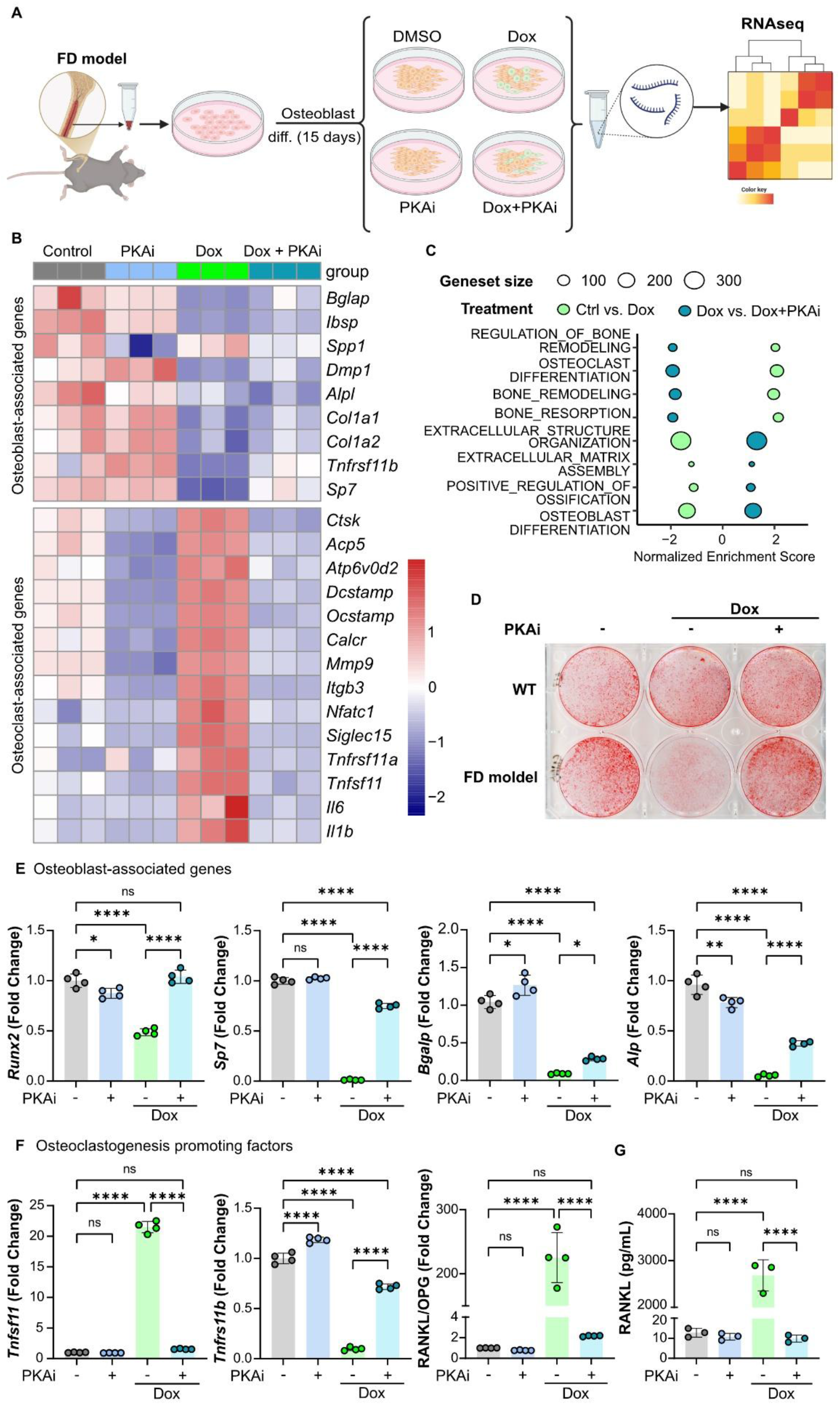
PKA inhibition restores osteoblast differentiation transcriptional programs and suppresses osteoclastogenic signaling induced by GNAS^R201C^. **A.** Experimental design. BMSCs isolated from FD mice were differentiated toward the osteoblast lineage for 15 days in the presence of vehicle (DMSO), doxycycline (Dox), PKA inhibitor (PKAi), or Dox+PKAi prior to RNA isolation and transcriptomic analysis by RNA sequencing. Created in BioRender. **B.** Heatmap showing expression of osteoblast-associated genes and osteoclast-associated genes across treatment conditions. **C.** Gene set enrichment analysis (GSEA) comparing Dox- and Dox+PKAi-treated osteoblast cultures. Bubble plot showing significantly enriched biological pathways. **D.** Representative Alizarin Red staining of wild-type (WT) and FD osteoblast cultures treated with Dox in the presence or absence of PKAi. Mineralized matrix deposition was assessed after osteoblast differentiation. Quantitative RT-PCR analysis of **E.** osteoblast-associated genes (*Runx2, Sp7, Bglap,* and *Alpl*) and **F.** osteoclastogenesis-promoting factors (*Tnfsf11* [RANKL], *Tnfrsf11b* [OPG], and the RANKL/OPG ratio) in osteoblast cultures treated with Dox in the presence or absence of PKAi. **G.** Quantification of secreted RANKL protein levels in conditioned media from osteoblast cultures treated with Dox and/or PKAi, as determined by ELISA. Data are presented as mean ± SD. Each dot represents an independent biological replicate. Statistical significance was determined by one-way ANOVA with multiple-comparison correction. ns, not significant; *P < 0.05, ****P < 0.0001.

Pathway analyses mirrored these gene-level changes. Signatures related to bone remodeling, osteoclast differentiation, and bone resorption were strongly activated under *GNAS*^R201C^ induction but were normalized following PKA inhibition. In addition, pathways associated with ossification and osteoblast differentiation, which were suppressed by Dox, were re-established in the presence of PKAi, indicating that inhibition of PKA catalytic activity promotes a shift back toward proper osteoblast identity and function (**Fig. 6C**).

These transcriptional changes translated into measurable improvements in osteoblast maturation. Alizarin Red staining showed a clear increase in calcium deposition in PKAi-treated cultures compared with Dox alone, demonstrating enhanced mineralization capacity and functional rescue of osteoblast maturation (**Fig. 6D**). qPCR analysis further confirmed the upregulation of osteoblast lineage factors (*Runx2* and *Sp7*) and mature osteoblast markers (*Alp* and *Bgalp*) following PKA inhibition (**Fig. 6E**).

Finally, we assessed whether PKAi impacts osteoblast-derived pro-osteoclastogenic factors. Doxycycline-induced cultures exhibited elevated expression of osteoclastogenic regulators, including increased secretion of RANKL into the conditioned media. Treatment with PKAi resulted in a marked reduction in these factors, including a significant decrease in RANKL secretion, thereby reducing the osteoblast-mediated drive toward osteoclastogenesis (**Fig. 6F-G**).

These findings demonstrate that PKA inhibition with BLU0588 restores osteoblast differentiation, enhances mineralization, suppresses aberrant osteoclastogenic signaling, and reactivates osteoblast-specific transcriptional pathways in *GNAS*^R201C^ BMSCs. This establishes PKA as a key mechanistic node whose pharmacologic blockade corrects both the molecular and functional defects in osteoblast maturation observed in FD.

### 6. PKA-mediated phosphorylation of RUNX2 suppresses osteolineage fate and enhances osteoclastogenic signaling

Considering the impact of PKA inhibition on osteoblast programs, we next sought to identify direct downstream PKA substrates that mediate osteolineage dysregulation. Through the analysis of multiple candidate osteogenic regulators we identified RUNX2 as a particularly compelling target, owing to both its central role as a master regulator of osteogenesis^15^ and the presence of putative PKA phosphorylation motifs. We therefore asked whether PKA directly acts on RUNX2 to disrupt its function and thereby impair osteoblast maturation.

To determine whether RUNX2 can undergo PKA-dependent phosphorylation, we performed RUNX2 isolation by pulldown assays in HEK293T cells expressing GST-RUNX2 and activating cAMP/PKA signaling with forskolin (stimulates adenylyl cyclase directly to increase the production of cAMP) and IBMX (inhibits the breakdown of cAMP) (F/I). Immunodetection assays using a PKA substrate antibody revealed a clear phosphorylation signal at the expected molecular weight of isolated GST-RUNX2, indicating that PKA stimulation promotes the phosphorylation of RUNX2. We found that PKAi abolished this signal, confirming PKA-dependency on RUNX2 phosphorylation (**Fig. 7A**). Guided by motif prediction (Scansite 4.0), we evaluated five candidate serine residues within the RUNX2 primary sequence located in the context of PKA-phosphorylation motifs (S28, S196, S198, S269, and S340) (**Supplementary Fig. 2A-B**). Systematic evaluation of mutated sites revealed S340 as the primary phosphorylation site in RUNX2 stimulated by PKA activity, and alanine substitution at this site substantially reduces the specific PKA substrate phosphorylation signal (**Fig. 7B**).

**Figure 7.**
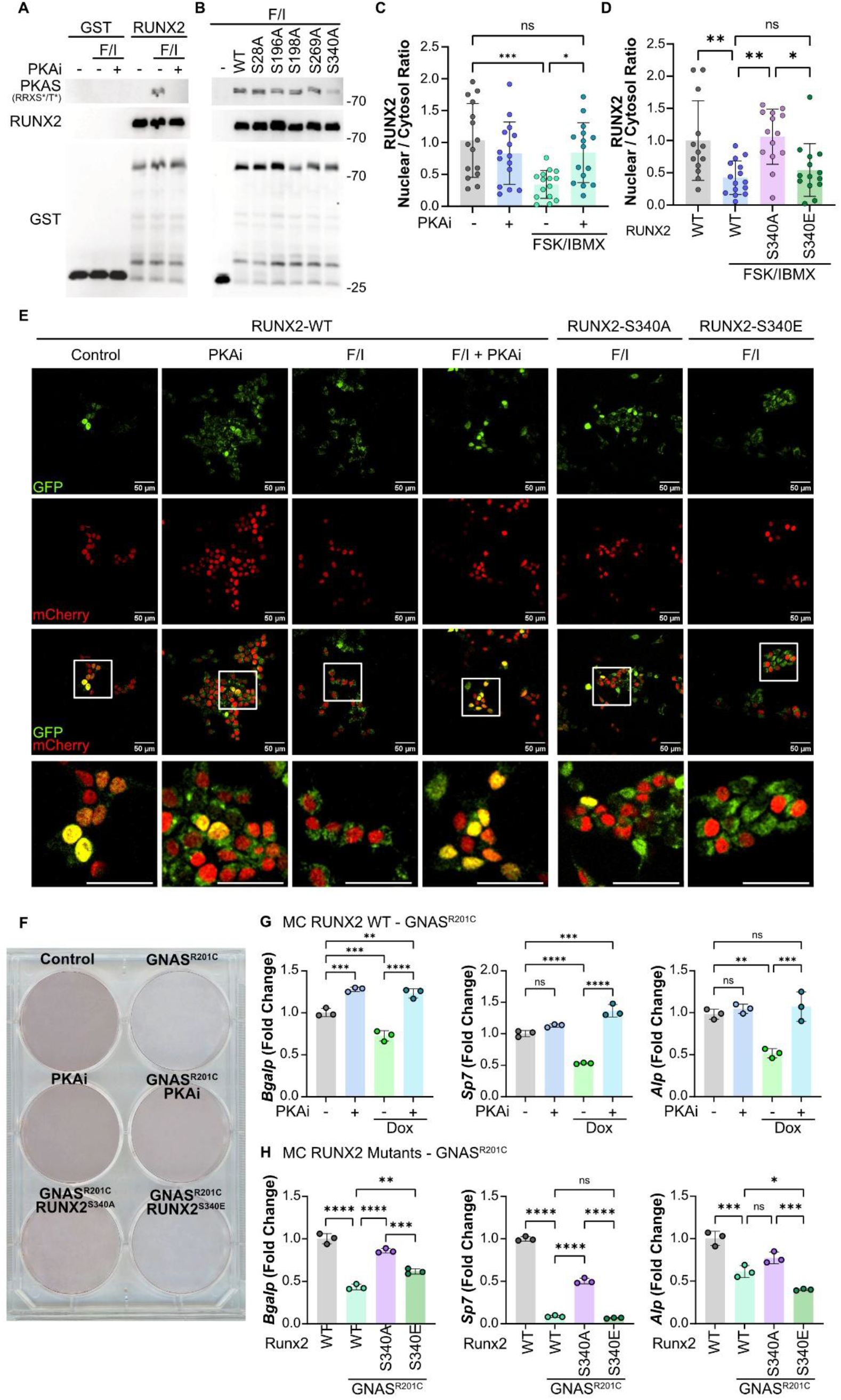
PKA stimulates the phosphorylation of RUNX2, thereby impairing osteoblast differentiation downstream of GNAS^R201C^. **A.** HEK293T cells were transfected with GST or GST-RUNX2 expression plasmids and treated with forskolin/IBMX (F/I) in the presence or absence of a PKA inhibitor (PKAi). GST pull-downs were analyzed by immunoblotting for phosphorylated PKA substrates (RRXS/T*) and RUNX2. **B.** HEK293T cells were transfected with GST-tagged RUNX2 wild-type (WT) or the indicated RUNX2 phospho-site mutants (S28A, S96A, S198A, S269A, and S340A). Following forskolin/IBMX (F/I) treatment, GST pull-downs were analyzed by immunoblotting for phosphorylated PKA substrates (RRXS/T*) and RUNX2. **C.** Quantification of the nuclear-to-cytosolic RUNX2 ratio in HEK293T cells treated with forskolin/IBMX (FSK/IBMX) in the presence or absence of PKAi. **D.** Quantification of the nuclear-to-cytosolic ratio of RUNX2-WT, RUNX2-S340A, and RUNX2-S340E in HEK293T cells following FSK/IBMX treatment. **E.** Representative fluorescence microscopy images of HEK293T cells expressing GFP-tagged RUNX2 constructs (green) together with H2B mCherry (red). Cells expressing RUNX2-WT, RUNX2-S340A, or RUNX2-S340E were treated with vehicle, PKAi, FSK/IBMX, or FSK/IBMX+PKAi as indicated. Insets show higher magnification views. Scale bars, 50 μm. **F.** Representative Alizarin Red staining of MC3T3-E1 subclone 4 cells (MC) expressing GNAS^R201C^, RUNX2-S340A, or RUNX2-S340E, as indicated. Mineralized matrix deposition was assessed following osteoblast differentiation. **G.** Quantitative RT-PCR analysis of osteoblast-associated genes (*Bglap, Sp7,* and *Alpl*) in MC3T3-E1 subclone 4 cells expressing RUNX2-WT and GNAS^R201C^ in the presence or absence of PKAi. **H.** Quantitative RT-PCR analysis of osteoblast-associated genes (*Bglap, Sp7,* and *Alpl*) in MC3T3-E1 subclone 4 cells expressing GNAS^R201C^ together with RUNX2-WT, RUNX2-S340A, or RUNX2-S340E. Data are presented as mean ± SD. Each dot represents an independent biological replicate. One-way ANOVA determined statistical significance with multiple-comparison correction. ns, not significant; *P < 0.05, **P < 0.01, ***P < 0.001, ****P < 0.0001.

We next asked how PKA-dependent phosphorylation influences RUNX2 localization. Using a RUNX2-GFP fusion protein and immunofluorescence microscopy, we found that acute activation of the PKA pathway maintained RUNX2 in the cytosol (**Fig. 7C**), reducing its nuclear accumulation. In contrast, the non-phosphorylatable S340A mutant remained predominantly nuclear under the same conditions, suggesting that phosphorylation at S340 is required for PKA-mediated nuclear exclusion. Consistently, a phosphomimetic mutant (S340E) showed constitutive cytosolic enrichment even in the absence of PKA activation, suggesting a direct role of S340 phosphorylation in RUNX2 cytosolic sequestration (**Fig. 7D**).

We validated these regulatory effects in an osteogenic cellular context using MC3T3-E1 subclone 4 (MC) cells, a widely used pre-osteoblast experimental model system that recapitulates key aspects of osteoblast differentiation and mineralization *in vitro*^16^. As a control, treatment of MC cells with cholera toxin, which persistently activates Gαs^17^ for 15 days, reduced mature osteoblast marker expression and markedly decreased Alizarin Red staining. A similar phenotype was observed following CRISPR-Cas9 sgRNA-mediated knockout of RUNX2, underscoring its essential role in osteoblast differentiation and supporting impaired maturation in both conditions (**Supplementary Fig 3A-C**). Consistent with these findings, overactivation of PKA (via expression of *GNAS*^R201C^) in MC cells impaired osteoblast differentiation and increased osteoclastogenic factor expression, recapitulating the osteolineage imbalance observed in our BMSC system (**Fig. 7F-G**). To validate the model, **Supplementary Fig. 4A-B** show efficient expression of the inducible *GNAS* mutant and activation of downstream PKA signaling, confirming the integrity of the experimental system.

We next performed rescue experiments in CRISPR-generated RUNX2 KO MC cells. Re-expression of wild-type RUNX2 fully restored osteogenic capacity, as evidenced by the recovery of mature osteoblast marker expression, indicating efficient rescue of the differentiation defect (**Supplementary Fig. 4C-E**). In this context, expression of the phospho-deficient RUNX2-S340A mutant similarly restored osteogenic differentiation and mineralization while reducing osteoclastogenic factor production. In contrast, the phosphomimetic RUNX2-S340E mutant failed to rescue differentiation, supporting a model in which PKA-mediated phosphorylation of RUNX2 impairs its pro-osteogenic function. (**Fig. 7F and H**).

Our results suggest that PKA stimulates the phosphorylation of RUNX2 at S340, controlling RUNX2 nuclear-cytoplasmic shuttling. Together, our findings offer a mechanistic link between PKA-stimulated retention of RUNX2 in the cytoplasm and impaired osteogenic transcription, suggesting that blocking RUNX2 S340 phosphorylation could restore its transcriptional function and rescue osteoblast differentiation even under conditions of *GNAS*-PKA hyperactivation.

### 7. PKA inhibition reduces lesion burden and restores bone structure *in vivo*

To evaluate the therapeutic potential of PKA inhibition *in vivo*, we used our inducible FD mouse model and first allowed FD-like lesions to form by administering Dox for one week (confirmed by increased limb thickness, which correlated with inflammation). After this induction period, animals were divided into four groups: (1) control mice that did not develop lesions, (2) control mice that received PKAi (45 mg/kg), (3) FD control mice maintained under Dox to sustain lesion development, and (4) FD mice treated with the PKAi for two weeks while continuing Dox exposure. This design allowed us to assess whether PKA blockade could reverse established FD-like pathology and whether PKAi treatment would affect normal mice. Across multiple measurements, PKA inhibition markedly improved bone phenotype. Gross examination revealed that PKAi-treated animals exhibited a reduction in limb swelling and deformity compared with FD controls (**Fig. 8A, Supplementary Fig 5A**). Micro-CT analysis further demonstrated a significant increase in bone density in PKAi-treated animals, accompanied by visible recalcification of previously lytic lesions, indicating active structural repair of the bone matrix (**Fig. 8B-C**).

**Figure 8.**
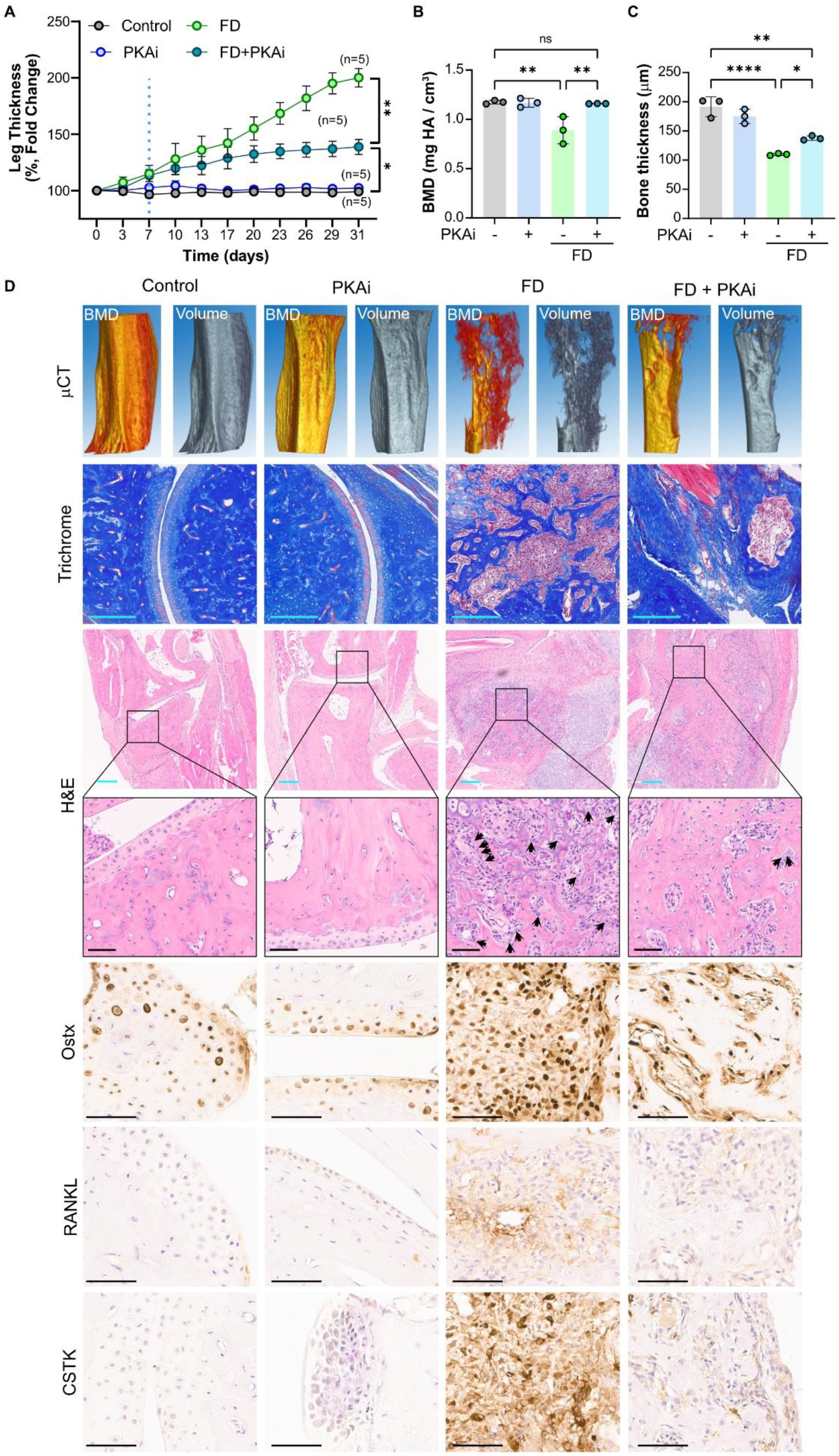
Pharmacological inhibition of PKA halts disease progression and reverses osteolytic pathology in GNAS^R201C^-driven FD mouse models. **A.** Longitudinal assessment of hindlimb thickness in control mice, FD mice, PKA inhibitor (PKAi)-treated control mice, and PKAi-treated FD mice. PKAi administration was initiated at day 7 (blue dotted line). Leg thickness is presented as percentage fold change relative to baseline measurements. **B.** Quantification of bone mineral density (BMD) and **C.** cortical bone thickness determined by micro-computed tomography (μCT) analysis of control, PKAi, FD, and FD+PKAi mice. **D.** Representative μCT reconstructions, Masson’s trichrome staining, hematoxylin and eosin (H&E) staining, and immunohistochemical analyses of Osterix (OSX), RANKL, and cathepsin K (CSTK) in femoral lesions from control, PKAi, FD, and FD+PKAi mice. Higher-magnification views of H&E-stained sections are shown. Arrows indicate osteoclast-rich areas within fibrous dysplasia lesions. Cyan scale bars represent 200 µm, and the black scale bars represent 60 µm. Data are presented as mean ± SD. Each dot represents an individual mouse. One-way ANOVA was used to determine statistical significance with multiple-comparison correction. ns, not significant; *P < 0.05, **P < 0.01, ****P < 0.0001.

Consistent with these findings, whole-mount skeletal staining demonstrated a marked improvement in femoral architecture and curvature following PKA inhibition, indicating partial correction of the gross skeletal abnormalities associated with FD (**Supplementary Fig 5B**). High-resolution MRI provided complementary confirmation of these structural improvements. FD control animals showed the characteristic increased porosity of FD-affected bone, whereas PKAi-treated animals showed a clear reduction in bone porosity, consistent with micro-CT findings. Quantitative MRI metrics supported this normalization: both the suppression ratio (bound water) and the magnetization transfer ratio (collagen) were increased in PKAi-treated mice, reflecting enhanced matrix integrity and improved bone tissue composition after treatment (**Supplementary Fig 5C-F**).

Finally, Masson’s trichrome staining revealed a marked reduction in fibrotic tissue inside bone following PKAi treatment. Hematoxylin and eosin (H&E) staining further showed fewer multinucleated cells, consistent with reduced numbers of mature osteoclasts. In line with these observations, PKAi treatment reduced osterix immunoreactivity, suggesting restored osteogenic activity. Additionally, RANKL expression was reduced, indicating decreased osteoclastogenic signaling, which correlated with a lower cathepsin K (CTSK) signal (**Fig. 8D**). Together, these findings demonstrate that pharmacological inhibition of PKA can reverse established FD lesions *in vivo* by restoring bone structure, reducing fibrotic remodeling, and normalizing osteoblast– osteoclast coupling, supporting PKA as a viable therapeutic target for disease modification in FD.

## Discussion

Our findings show how pathogenic GNAS mutations disrupt normal osteogenic programs in skeletal stem cells, demonstrate that pharmacological inhibition of *PRKACA* reverses established FD lesions *in vivo*, and identify a previously unrecognized molecular mechanism linking aberrant *GNAS*–PKA signaling to impaired osteoblast differentiation in FD. Specifically, we identified RUNX2, the master transcriptional regulator of osteogenesis^18^, as a target of PKA activity and demonstrate that phosphorylation at S340 acts as a negative regulatory switch that suppresses RUNX2 nuclear activity. Mechanistically, we provide evidence that this modification promotes cytoplasmic sequestration of RUNX2, inhibits osteogenic transcriptional programs, and impairs osteoblast maturation. The ability of the phospho-deficient RUNX2-S340A mutant to restore osteogenic differentiation despite persistent *GNAS*^R201C^ signaling supports RUNX2 as a critical downstream effector of PKA-driven FD pathology.

Of interest, structural analysis suggests that S340 lies within the proline/serine/threonine-rich (PST) domain of RUNX2, which contains activation domain 3 (AD3) and the nuclear matrix targeting signal (NMTS), both of which are required for proper nuclear localization and transcriptional activity^19, 20^. Disruption of the PST domain has been shown to alter RUNX2 subcellular distribution from a predominantly nuclear localization to a diffuse nucleo-cytoplasmic pattern^19^, underscoring the importance of this region in regulating RUNX2 function. Notably, the sequence surrounding S340 conforms to a canonical PKA consensus motif, and the conservation of this residue across all vertebrate species suggests an evolutionarily conserved regulatory role (Supplementary Fig 2C). Together with our findings, these observations support S340 as a candidate signaling node through which PKA may regulate RUNX2 activity.

Phosphorylation-dependent control of nuclear trafficking is a well-established mechanism by which signaling pathways regulate transcription factor function. PKA phosphorylation can inhibit nuclear accumulation either by reducing importin-mediated transport or by promoting cytoplasmic retention, as described for NFATc1^21^ and APC^22^. Alternatively, phosphoserine residues can recruit 14-3-3 proteins that mask nuclear localization signals and prevent nuclear import, as reported for FOXO family members and class IIa HDACs^23, 24^. Although the precise mechanism remains to be determined, our observation that the phosphomimetic S340E mutant is largely excluded from the nucleus and fails to rescue osteoblast differentiation suggests that phosphorylation at S340 is sufficient to impair RUNX2 nuclear function. These findings suggest that PKA-mediated phosphorylation of RUNX2 represents a previously unrecognized mechanism linking aberrant *GNAS*-initiated cAMP–PKA signaling to defective osteoblast differentiation in fibrous dysplasia.

We leveraged BLU0588, a recently developed selective inhibitor of PKA catalytic activity^8, 9^, to assess PKA pathway dependency and explore the therapeutic potential of PKA inhibition in this disease context. On-target inhibition was confirmed by suppression of canonical PKA targets (e.g., pVASP and CRE-reporter activity), without affecting upstream *GNAS* expression. Functionally, *PRKACA* blockade restored osteoblast differentiation programs and mineralization, while concomitantly reducing osteoclastogenic signaling and activity. These dual effects are consistent with the known role of osteoblast-lineage cells as key regulators of osteoclastogenesis through RANKL production^5, 25, 26^, which is elevated in FD lesions and drives excessive osteoclast formation. Notably, independent studies using genetic and pharmacologic strategies targeting the RANKL axis in *GNAS*-driven FD models have similarly shown reduced fibrosis, decreased osteoclast burden, and partial restoration of osteoblast differentiation^3, 5^. This supports the concept that modulation of the osteoblast–osteoclast crosstalk is sufficient to alter disease trajectory, which we can now achieve by targeting PKA, as a proximal *GNAS*-induced pathologic signaling.

From a translational perspective, our study highlights *PRKACA* inhibition as a disease-modifying strategy that, unlike anti-resorptive therapies, simultaneously addresses both arms of FD pathology: defective bone formation and excessive bone resorption. While agents such as denosumab effectively suppress osteoclast activity and can transiently improve bone architecture, their effects are often incomplete and reversible upon drug withdrawal^5, 6^. In contrast, targeting the upstream *GNAS*–PKA axis can reprogram osteoprogenitor cell fate, restore RUNX2 function, and rebalance the skeletal microenvironment. This dual mechanism of action may offer durable therapeutic benefit across disease stages and anatomical sites. Moreover, given the convergence of multiple signaling pathways on PKA, combination strategies that integrate PKA inhibition with RANKL blockade or other modulators of the bone microenvironment may further enhance structural repair and limit disease relapse.

In summary, our work identifies a mechanistic axis linking *GNAS* mutations to impaired osteoblast differentiation via PKA-dependent phosphorylation of RUNX2, and provides compelling preclinical evidence that pharmacologic inhibition of *PRKACA* can reverse FD pathology. These findings advance our understanding of disease biology and provide a strong rationale for targeting the *GNAS*–PKA–RUNX2 pathway in skeletal stem cells as a precision therapeutic strategy in FD.

## Limitation of the study

The bulk transcriptomic analyses used here do not fully capture the cellular heterogeneity and spatial organization of FD lesions, which are increasingly recognized as complex mosaics of mutant and wild-type cell populations interacting within a fibrotic niche. Recent single-cell studies of human FD tissue have revealed distinct stromal, endothelial, and perivascular populations that contribute to lesion architecture and fibrosis, highlighting the need for future higher-resolution approaches to refine the cellular targets of PKA inhibition^27^. Additionally, while S340 emerges as a dominant regulatory site on RUNX2 in our system, RUNX2 may integrate multiple upstream kinase inputs, and comprehensive phosphoproteomic analyses will be necessary to define the broader signaling network controlling its activity in FD. Finally, developing bone-specific pharmacodynamic biomarkers of PKA inhibition will be essential to guide and expedite clinical translation and optimize therapeutic dosing.

## Conclusion

Our findings uncover a previously unrecognized mechanism through which aberrant *GNAS*–PKA signaling disrupts skeletal stem cell osteogenic cell fate and drives FD pathogenesis. By identifying RUNX2 as a critical downstream effector of PKA and demonstrating the reversal of established lesions through pharmacological *PRKACA* inhibition, this study provides both a mechanistic framework for disease progression and a rationale for future targeted therapeutic intervention. Together, these results establish the *GNAS*–PKA–RUNX2 axis as a central regulator of skeletal homeostasis and a promising therapeutic vulnerability in FD and other bone-related diseases.

## Authors contribution

Conceptualization, E.S.C-A and J.S.G.; methodology, E.S.C-A., D.R., A.S., T.I., S.R.A-G., T.O., and S.H.S., validation E.S.C-A., D.R., A.S., K.S., T.I., and J.K.; formal analysis E.S.C-A, K.S. and S.H.S, investigation E.S.C-A., S.R.A-G.; resources J.S.G., and F.F.; data curation E.S.C-A., and K.S.; original draft preparation E.S.C-A.; writing reviewing and editing E.S.C-A and J.S.G., visualization E.S.C-A. and J.S.G; supervision J.S.G., F.F. and J.D.; project administration E.S.C-A., and J.S.G.; funding acquisition E.S.C-A and J.S.G. All authors have read and agreed to the published version of the manuscript.

## Acknowledgments

Part of this project was funded thanks to FD/MAS Alliance and Team FD/MAS in the UPenn Million Dollar Bike Ride.

## Extended data figures

Extended data figures 1–5

## Online Methods

### Cell culture

HEK293T (ATCC, Cat# CRL-3216) and MC3T3-E1 subclone 4 (ATCC, Cat# CRL-2593) cells were obtained from ATCC. HEK293A cells (Cat# R70507) were purchased from Invitrogen. Cells were maintained for no more than 15 passages. Cells were routinely tested for mycoplasma contamination monthly using a Mycoplasma Detection Kit (MedChem Express, Cat# HY-K0552).

HEK293T and HEK293A cells were cultured in Dulbecco’s Modified Eagle Medium (DMEM; Gibco, Cat# 11965118) supplemented with 10% fetal bovine serum (FBS; Sigma-Aldrich, Cat# 12103C), 1× MEM non-essential amino acids (Gibco, Cat# 11140-050), and 1× antibiotic-antimycotic solution (Gibco, Cat# 15240-062). MC3T3-E1 subclone 4 cells (hereafter referred to as MC cells) were cultured in α-MEM without ascorbic acid (Gibco, Cat# A10490-01) supplemented with 10% FBS and 1× antibiotic-antimycotic solution (complete medium). Cells were maintained at 37 °C in a humidified incubator with 5% CO₂.

For cryopreservation, cells were resuspended in freezing medium consisting of 80% FBS, 10% dimethyl sulfoxide (DMSO; Sigma-Aldrich, Cat# D8418), and 10% complete medium. Cells were frozen at −80 °C for 24 h using a controlled-rate freezing container (Mr. Frosty) and subsequently transferred to liquid nitrogen for long-term storage.

### Fibrous dysplasia animal model

All animal studies were approved by the University of California San Diego Institutional Animal Care and Use Committee (IACUC; protocol number S15195) and conducted in accordance with institutional and federal guidelines. Transgenic mouse strains were bred and maintained as previously described^4^. Control mice (*Prrx1*-Cre; Rosa26^LSL-rtTA^; TetO-H2B-GFP) and fibrous dysplasia (FD) model mice (*Prrx1*-Cre; TetO-Gsα^R201C^; Rosa26^LSL-rtTA^; TetO-H2B-GFP) were used for all in vivo experiments. Doxycycline (0.05 g/L) was administered in the drinking water starting at 7 weeks of age to induce transgene expression, and baseline limb thickness was recorded at the time of induction. Detectable limb inflammation developed in FD mice within 7 days, after which animals were randomized into treatment groups (n = 5 per group, males and females). Control mice were assigned to vehicle or PKA inhibitor (PKAi; BLU0588), and FD mice were assigned to parallel treatment groups (vehicle or FD + PKAi). PKAi was administered once daily by oral gavage at 45 mg/kg. Limb thickness was measured every 72 hours, and animals were maintained on treatment for 3 additional weeks. For pain management, all mice received carprofen (5 mg/kg) once daily starting at group randomization. At the experimental endpoint, animals were euthanized, and mandibles and limbs were collected. Hindlimbs were fixed in Zinc Formalin Fixative (Z-fix) (Sigma-Aldrich, Cat# Z2902) for 48 hours and then transferred to 70% ethanol for storage and downstream processing. One limb per animal was preserved intact for MRI and µCT imaging.

### Histology and Immunohistochemistry

Harvested bones were fixed in Z-fix, decalcified with StatLab™ Immunocal Decalcifier (StatLab, Cat# STL141432), dehydrated through graded alcohols, and embedded in paraffin. Sections (5 µm) were prepared and stained with Hematoxylin and eosin (H&E), Masson’s trichrome, or subjected to immunohistochemistry (IHC). IHC was performed using the following primary antibodies: anti-Cathepsin K (E7U5N, 1:400, Cell Signaling Technology, Cat# 57056), anti-RANKL (12A668, 1:200, Abcam, Cat# ab45039), and anti-Sp7/Osterix (polyclonal, 1:200, Abcam, Cat# ab22552). The bound antibodies were detected using Goat anti-Rabbit IgG Antibody (H+L), Biotinylated (1:400, Vector laboratories, Cat# BA-1000) or Goat Anti-Mouse IgG Antibody (H+L), Biotinylated (1:400, Vector laboratories, Cat# BA-9200), followed by the VECTASTAIN® Elite ABC-HRP Kit, Peroxidase (Standard) (Vector laboratories, Cat# PK-6100) and ImmPACT® DAB Substrate, Peroxidase (HRP) (Vector laboratories, Cat# SK-4105), according to the manufacturer’s instructions. Stained slides were scanned using an Aperio AT2 microscope slide scanner (Leica) for subsequent analysis. Appropriate positive and negative controls were included in each staining run.

### Whole skeletal staining

Whole-mount skeletal staining was performed as previously described^4^. Briefly, mice were euthanized and eviscerated, and skin and soft tissues were removed before fixation with 95% ethanol. Skeletal preparations were stained with Alcian Blue to visualize cartilage and Alizarin Red to label mineralized bone, followed by clearing in graded glycerol solutions. Stained specimens were imaged using standard stereomicroscopy.

### Bone marrow stromal cell (BMSC) isolation

Femurs and tibias were isolated from 7-week-old FD mice and placed in ice-cold PBS. Under sterile conditions, the epiphyses were removed, and bone marrow was collected by brief centrifugation (15 s at 9,000 × g). Bones were discarded, and the marrow pellet was resuspended and dissociated using a 23-gauge needle. Cells were resuspended in complete medium consisting of α-MEM supplemented with GlutaMAX (Gibco, Cat# 32571036), 20% fetal bovine serum, and 1× antibiotic-antimycotic solution, passed through a 70 µm cell strainer, counted, and plated at a density of 5 × 10⁶ cells per 10 cm dish. After 4 days, non-adherent cells were removed by washing with PBS, and fresh complete medium was added. Cultures were maintained with medium changes every 2 days and used for experiments 10 days after isolation.

### Osteogenic and osteoclastogenic differentiation

BMSCs were seeded at a density of 5 × 10⁵ cells per well in 6-well plates and allowed to reach ∼80% confluence. Upon attachment, the medium was replaced with either osteogenic or osteoclastogenic differentiation medium. Osteogenic differentiation was induced using α-MEM supplemented with GlutaMAX, 10% fetal bovine serum, 10 mM β-glycerol phosphate (Sigma, Cat# G9422), 200 µM ascorbic acid (Cayman, Cat# 14656), and 100 nM dexamethasone (Cayman, Cat#11015)^10^, and cells were maintained for 15 days. Complete differentiation was confirmed by alizarin red staining (Cosmo Bio Co, Cat# CSR-ARD-A1) according to the manufacturer’s instructions. For MC cells, differentiation medium consists of complete medium supplemented with L-ascorbic acid (50 µg/mL) and β-glycerophosphate (10 mM)^28^. Cells were differentiated for 21 days with media changes every 3 days.

Osteoclast differentiation was induced using α-MEM supplemented with GlutaMAX, 10% fetal bovine serum, 25 µg/mL RANKL (Thermo Fisher Scientific, Cat# 310-01), and 25 µg/mL M-CSF (Thermo Fisher Scientific, Cat# 315-02)^11^, and cells were maintained for 7 days. Medium was replaced every 3 days throughout the differentiation period. Differentiation was assessed using the TRAP kit (Sigma, Cat# 387A-1KT) according to the manufacturer’s instructions.

### RNA isolation and quantitative PCR

Total RNA was isolated using the Monarch® Total RNA Miniprep Kit (New England Biolabs, Cat# T2110L) according to the manufacturer’s instructions. RNA concentration and purity were determined using a NanoDrop spectrophotometer. For cDNA synthesis, 1 µg of total RNA was reverse-transcribed using Luna® Reverse Transcriptase (New England Biolabs, Cat# E3010) following the manufacturer’s protocol. The resulting cDNA was diluted 1:50 before quantitative PCR analysis. Quantitative PCR (qPCR) was performed using PowerTrack™ SYBR Green Master Mix (Thermo Fisher Scientific, Cat# A46109) on a CFX384 Real-Time PCR Detection System (Bio-Rad). Reactions were carried out under standard cycling conditions recommended by the manufacturer. Relative gene expression was calculated using the 2^−ΔΔCt method, with GAPDH used as the internal control, and results are presented as fold change relative to the control condition. Primer sequences are listed in Table 1.

**Table 1.**
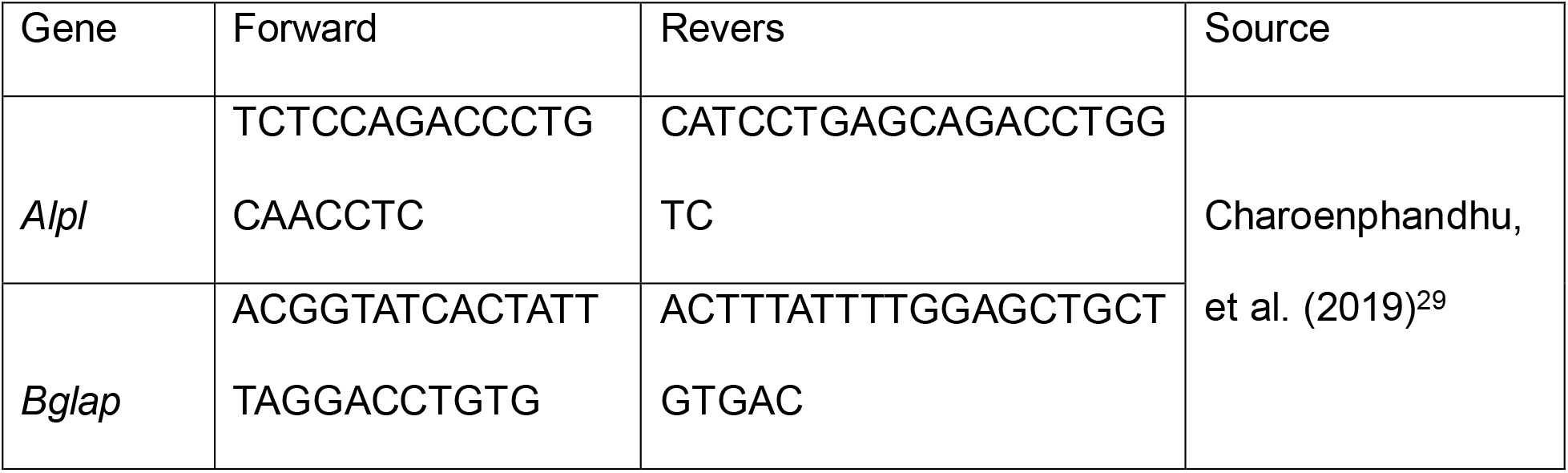
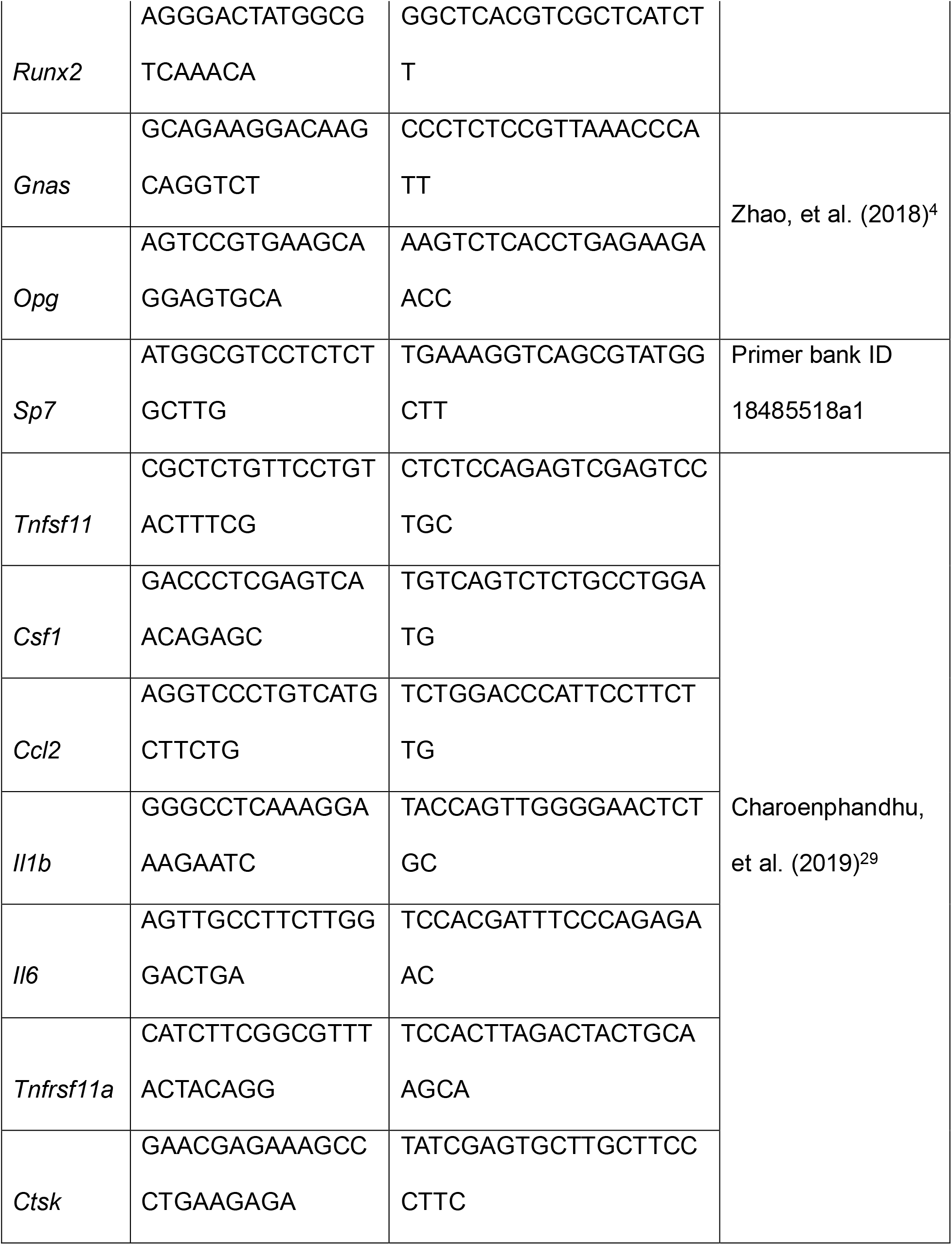
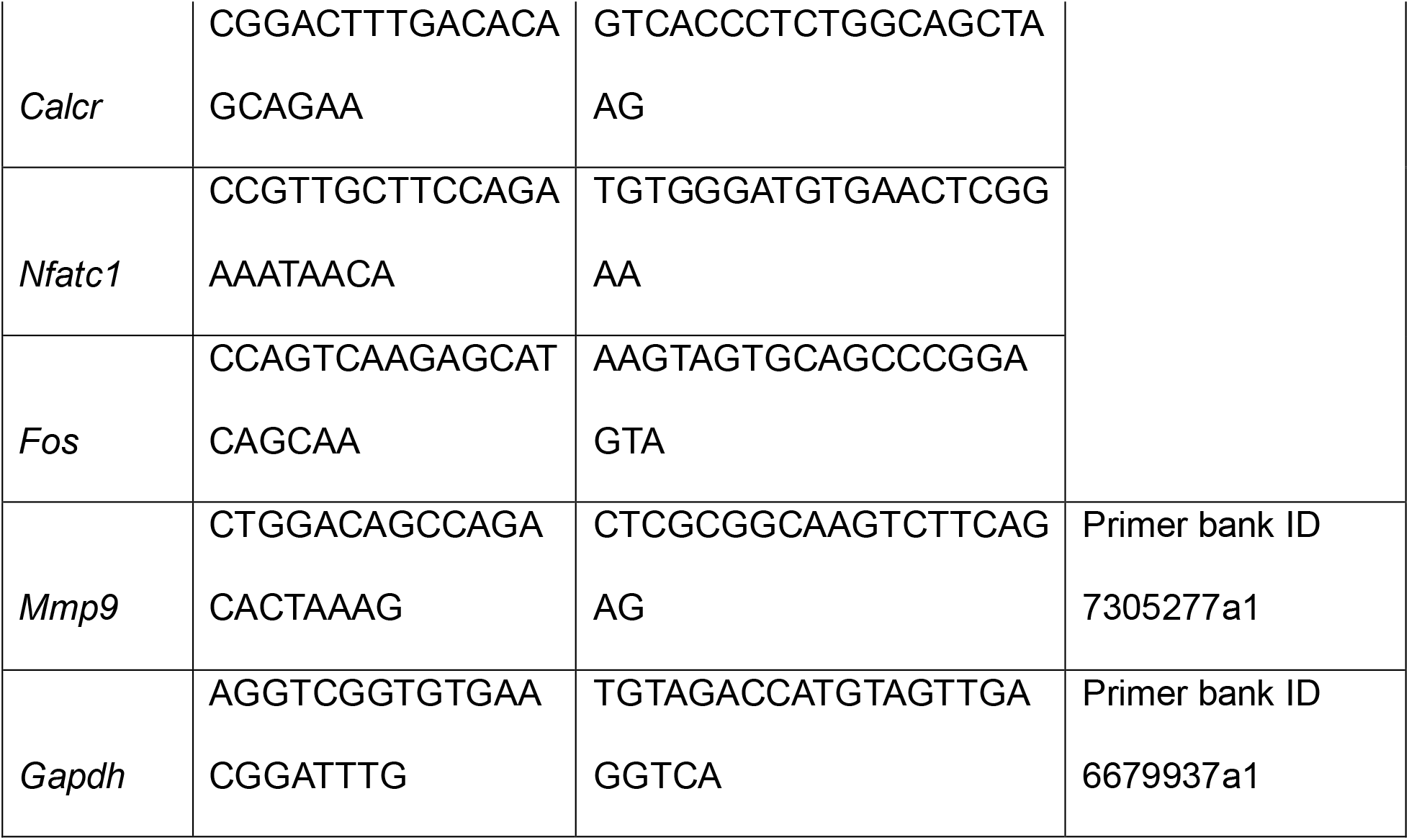
qPCR primers Mouse.

### siRNA and transfection

Cells were transfected with 20 nM of SMARTpool siRNAs (Horizon Discovery) using Lipofectamine RNAiMAX reagent (Invitrogen, Cat# 13778150) according to the manufacturer’s instructions. Culture media were refreshed 24 h after transfection. Cells were incubated with the corresponding treatments for 48 h and collected for processing.

### Protein extraction

Cells were lysed in ice-cold RIPA buffer (Thermo Fisher Scientific, Cat# 89900) supplemented with protease and phosphatase inhibitors (Thermo Fisher Scientific, Cat# 78447) and sodium orthovanadate (New England Biolabs, Cat# P0758L). Lysates were incubated on ice for 5 min and then sonicated (Qsonica sonicator, 3 pulses of 5 s each at 25% amplitude). Samples were then centrifuged at 21,200 × g for 10 min at 4 °C to remove insoluble debris. The clarified supernatants were collected, and protein concentrations were determined using the Bio-Rad DC Protein Assay (Bio-Rad, Cat# 5000111). We used 25 µg of protein per sample per well for immunoblotting.

### GST pull-down assays

For GST pull-down experiments, cells transfected with GST-RUNX2 wild-type, mutants, or control plasmid were lysed in ice-cold IP lysis buffer (Thermo Fisher Scientific, Cat# 87788) supplemented with protease and phosphatase inhibitors (Thermo Fisher Scientific, Cat# 78447) and sodium orthovanadate (New England Biolabs, Cat# P0758L; 1:100 dilution) 48 hours post-transfection. Lysates were incubated on ice for 5 min, then sonicated using a Qsonica sonicator (4 pulses of 1 s each). Samples were centrifuged at 21,200 × g for 10 min at 4 °C to remove insoluble debris. Protein concentrations were determined using the Bio-Rad DC Protein Assay (Bio-Rad, Cat# 5000111), and all samples were normalized to equal protein amounts. An aliquot (50 µL) was collected from each lysate as an INPUT control.

Clarified lysates were incubated with glutathione Sepharose 4B beads (Sigma-Aldrich, Cat# GE17-0756-01) according to the manufacturer’s recommendations. Briefly, lysates were incubated with equilibrated beads under gentle rotation at 4 °C for 2 h to allow binding of GST-tagged proteins. Following incubation, beads were washed four times with IP lysis buffer to remove nonspecific interactions. Bound proteins were eluted by boiling in SDS sample buffer and subjected to immunoblot analysis.

### Immunoblotting

Proteins were resolved by SDS-PAGE using polyacrylamide gels of varying concentrations depending on the molecular weight of the target proteins and transferred to membranes using the Trans-Blot Turbo transfer system (Bio-Rad) with standard program settings. Membranes were blocked with 5% non-fat milk and incubated with primary antibodies followed by HRP-conjugated secondary antibodies according to standard protocols. The following primary antibodies were used: HSP90 (Cat# 4877S), Phospho-PKA Substrate (RRXS*/T*) (100G7E) (1:1000, Cat# 9624), VASP (1:1000, Cat# 3132S), phospho-VASP (Ser157) (1:1000, Cat# 3111S), GFP (1:5000, Cat# 2956), PKA catalytic α (1:1000, Cat# 4782), GST (1:5000, Cat# 2624), CREB (1:500, Cat# 9704S), phospho-CREB (1:1000, Cat# 9198S), and PKA RI-α/β (1:1000, Cat# 3927) (all from Cell Signaling Technology); Gαs (1:1000, Sigma-Aldrich, Cat# 371732); and RUNX2 (1:1000, Assay Biotechnology, Cat# R12-2340). Secondary antibodies used were anti-rabbit (1:20000, SouthernBiotech, Cat# 4010-05) and anti-mouse (1:20000, SouthernBiotech, Cat# 1010-05). Immunoreactive bands were detected using Pierce ECL Western Blotting Substrate (Thermo Fisher Scientific, Cat# 32106).

### RNA sequencing (RNA-seq) and bioinformatics analysis

Total RNA extracted from BMSCs, osteoblasts, and osteoclasts was subjected to paired-end 150bp sequencing on the Illumina NovaSeq6000 sequencer at Novogene Co., Ltd. Sequencing reads were aligned to GRCm39 mouse reference genome by STAR v2.7.9 using Gencode vM34 annotations. Read count data was generated by RSEM and differentially expressed genes were obtained by DESeq2. Gene set enrichment analysis (GSEA) was performed using the Mouse Molecular Signatures Database (MSigDB) v2024.1^30^. Gene ontology (GO) analysis was performed using Metascape^31^. Transcription factor (TF) activities were inferred using R package decoupleR v2.9.1^32^ and CollecTRI TF database ^33^.

### RANKL ELISA

RANKL levels in cell culture supernatants were measured using a mouse TRANCE/RANKL ELISA kit (R&D Systems, Cat# MTR00) according to the manufacturer’s instructions. Briefly, clarified supernatants and standards were added in triplicate to antibody-coated 96-well plates and incubated for 2 h at room temperature, followed by washing. HRP-conjugated detection antibody was added for 2 h, and signal was developed using TMB substrate. The reaction was stopped, and absorbance was read at 450 nm with wavelength correction at 540/570 nm. RANKL concentrations were calculated from a standard curve.

### Bone resorption assay

Osteoclast activity was assessed using a bone resorption assay kit (Cosmo Bio Co., Cat# CSR-BRA-24KIT) according to the manufacturer’s instructions. Briefly, cells were cultured on calcium phosphate–coated plates preloaded with fluorescein amine-labeled chondroitin sulfate under differentiation conditions for 7 days. Osteoclast-mediated resorption releases the fluorescent substrate into the culture medium. At the endpoint, supernatants were collected, and fluorescence was measured using a Tecan Spark multimode microplate reader (excitation 485 nm, emission 535 nm). Fluorescence intensity was used as a quantitative measure of bone resorption activity.

### CRE reporter assay

Cells were seeded in 12-well plates at ∼70% confluence. The following day, cells were co-transfected with a CRE reporter plasmid and a Renilla luciferase plasmid as an internal control using TurboFect Transfection Reagent (Thermo Fisher Scientific, Cat# R0531). Cells were maintained in transfection medium for 48 h, after which the medium was replaced with starvation medium, and cells were stimulated under the indicated conditions and time points. Luciferase activity was measured using the Dual-Glo Luciferase Assay System (Promega, Cat# E2920) according to the manufacturer’s instructions. Firefly luciferase activity was normalized to Renilla luciferase activity.

### Plasmids and site-directed mutagenesis

pCEFL GST backbone was obtained from a previously published construct^34^. The vector was linearized by digestion with BamHI (New England Biolabs, Cat# R0136S) downstream of the GST coding sequence, followed by dephosphorylation using recombinant Shrimp Alkaline Phosphatase (rSAP; New England Biolabs, Cat# M0371S). To generate GST–RUNX2 construct, the RUNX2 coding sequence was amplified from pPB-cT3G-cERP2-RUNX2, a gift from George Church^35^ (Addgene plasmid # 192926; http://n2t.net/addgene:192926;RRID:Addgene_192926), using gene-specific primers containing homology arms compatible with the BamHI-digested pCEFL GST vector. PCR amplification was performed using PrimeSTAR GXL DNA Polymerase (Takara, Cat# R050A). The parental template plasmid was digested with DpnI (New England Biolabs, Cat# R0176S) to remove methylated DNA. The amplified RUNX2 insert was cloned into the linearized vector using the NEBuilder HiFi DNA Assembly Master Mix (New England Biolabs, Cat# E2621S). Assembled plasmids were transformed, amplified, and validated by Sanger sequencing (Plasmidsaurus).

For GFP-tagged constructs, the same cloning strategy was used with the pCEFL GFP backbone^36^, which contains a BamHI cloning site analogous to the GST vector, to generate RUNX2–GFP fusion constructs.

Site-directed mutagenesis was performed on the pCEFL GST–RUNX2 plasmid using the QuikChange II Site-Directed Mutagenesis Kit (Agilent, Cat# 200523) to generate a panel of six mutants. Two of the mutations (340A and 340E) were also introduced into the RUNX2–GFP constructs using the same approach. For lentiviral expression, RUNX2 constructs were generated using pFUW-TetO-RUNX2, a gift from Filipe Pereira^37^ (Addgene plasmid # 207225; http://n2t.net/addgene:207225;RRID:Addgene_207225), as a backbone, and mutants were introduced using the QuikChange II XL Site-Directed Mutagenesis Kit (Agilent, Cat# 200521).

### CRISPR Knock Out RUNX2

RUNX2 knockout (KO) MC3T3-E1 cells were generated using CRISPR/Cas9 ribonucleoprotein (RNP) complexes (IDT, Cat# 1081058) and three guide RNAs. Cells were electroporated using a Neon Transfection System (1200 V, 10 ms, 3 pulses), and KO efficiency was validated by Western blot and qPCR.

### Lentivirus generation

Lentiviral particles were produced in HEK293T cells by co-transfection with packaging plasmids encoding VSV-G and psPAX2 at a ratio of 1:2:3 (VSV-G:psPAX2:pLenti plasmid). Viral supernatants were collected and concentrated using Lenti-X Concentrator (Takara Bio, Cat# 631231) according to the manufacturer’s instructions.

### Lentiviral Transduction and RUNX2 Reconstitution

MC3T3-E1 subclone 4 cells (MC cells), including WT and RUNX2 knockout (RUNX2 KO) populations, were transduced with a lentiviral vector encoding the constitutively active *GNAS*^R201C^ mutant (pLVX-TetOne *GNAS*^R201C^ FLAG ^38^) carrying puromycin resistance. Following infection, cells were selected with puromycin to establish stable cell populations.

For RUNX2 reconstitution experiments, MC RUNX2 KO cells were transduced with lentiviral particles encoding wild-type RUNX2 (pFUW-TetO-RUNX2) or RUNX2 mutants (S340A and S340E). As these vectors do not contain a selectable marker, transduced cells were used as pooled populations for downstream assays.

### Immunofluorescence imaging

HEK293T cells stably expressing mCherry-H2B were seeded onto glass-bottom culture dishes (MatTek, Cat# CCS-8) pre-coated with poly-D-lysine (Sigma-Aldrich, Cat# P6407) at ∼70% confluence. Cells were transfected with pCEFL-GFP-RUNX2 (wild-type or mutant constructs) using TurboFect transfection reagent (Thermofisher, Cat# R0531) according to the manufacturer’s instructions and maintained for 48 hours. Before stimulation, cells were serum-starved for 6 hours and then treated with the indicated stimuli. Cells were fixed with 4% paraformaldehyde (PFA) for 10 minutes at room temperature. Images were acquired using a confocal microscope (Nikon A1R Confocal). For image analysis, nuclear and cytoplasmic fluorescence intensities were quantified using Fiji (ImageJ). Nuclei were defined based on the mCherry-H2B signal, and corresponding regions of interest (ROIs) were generated. Cytoplasmic ROIs were defined by excluding the nuclear region from the total cell area. Mean fluorescence intensities for GFP-RUNX2 were measured within nuclear and cytoplasmic compartments, and nuclear-to-cytoplasmic ratios were calculated for each cell. Quantification was performed on multiple fields of view per condition.

### µCT and MRI Analysis

Samples were fixed in Z-Fix and subsequently stored in 70% ethanol before imaging. Micro-computed tomography (µCT) and magnetic resonance imaging (MRI) were performed to assess bone structure, mineralization, and composition.

For µCT analysis, bone specimens were imaged using a micro-computed tomography scanner (Skyscan 1174, Bruker, Kontich, Belgium) with an isotropic voxel size of 10.8 µm. Scans were performed in the presence of hydroxyapatite calibration phantoms (0.25 and 0.75 g/cm³) to quantify volumetric bone mineral density (vBMD). Acquisition parameters were 50 kVp, 0.8 mA, 2.2 s exposure time, 0.3° rotation step, and two-frame averaging, with a 0.25-mm aluminum filter to reduce low-energy X-ray artifacts. Local vBMD values were calculated by linear interpolation of grayscale intensities relative to the calibration phantoms. A threshold of 0.5 g/cm³ was applied to exclude non-bone regions prior to calculation of mean vBMD for each specimen.

Ultrashort echo time (UTE) MRI data were acquired on a Bruker 3T scanner using a surface cryoprobe coil^39^. Proton density (PD)-weighted dual-echo UTE images were obtained to calculate the porosity index (TR = 20 ms; TE1/TE2 = 0.016/2.2 ms; flip angle (FA) = 4°; number of averages (NA) = 2; scan time = 38 min). Short TR adiabatic inversion recovery UTE (STAIR-UTE) data were acquired to evaluate suppression ratio (TR/TI = 150/63 ms; TE = 0.019 ms; FA = 35°; NA = 10; inter-spoke TR = 4 ms; 9 spokes per inversion recovery pulse; scan time = 2 h 24 min). Magnetization transfer (MT)-weighted UTE (UTE-MT) images were acquired to assess matrix composition (TR/TE = 86/0.0135 ms; FA = 7°; NA = 6; inter-spoke TR = 7 ms; 9 spokes per MT pulse; scan time = 48 min). The MT pulse consisted of a Fermi-shaped pulse (flip angle = 1500°; offset frequency = 2000 Hz), and reference images (MT_off) were acquired with the MT pulse disabled. All MRI datasets were acquired with identical spatial parameters (matrix = 128 × 128 × 128; field of view (FOV) = 10 × 10 × 50 mm).

Parametric maps were calculated as follows: porosity index = PD-UTE TE2 / TE1; suppression ratio = STAIR-UTE / PD-UTE TE1; and magnetization transfer ratio (MTR) = (UTE-MT_off – UTE-MT_on) / UTE-MT_off.

## Statistics

Statistical analyses were performed using GraphPad Prism software. For comparisons between two groups, an unpaired two-tailed Student’s *t*-test was used. For comparisons among three or more groups, one-way ANOVA followed by Tukey’s post hoc test was applied when the data were normally distributed. For non-normally distributed data, nonparametric tests (e.g., Mann–Whitney test) were used as appropriate. Two-way ANOVA followed by Tukey’s multiple comparisons test was used to assess the effects of two independent variables simultaneously. Data are presented as mean ± standard error of the mean (SEM) from at least three independent experiments. Differences were considered statistically significant when *p* < 0.05. Statistical significance is indicated as follows: p < 0.05 (*), p < 0.01 (**), p < 0.001 (***), and p < 0.0001 (****).

## Declaration of generative AI and AI-assisted technologies in the writing process

During the preparation of this work, the authors used Copilot® and Grammarly® to check syntax and grammar, and appropriate text structure. After using these tools, the authors reviewed and edited the content as needed and took full responsibility for the content of the published article.

## Reference

1. Boyce, A.M. & Collins, M.T. Fibrous Dysplasia/McCune-Albright Syndrome: A Rare, Mosaic Disease of Galpha s Activation. Endocr Rev 41, 345–370 (2020).

2. Weinstein, L.S. et al. Activating mutations of the stimulatory G protein in the McCune-Albright syndrome. N Engl J Med 325, 1688–1695 (1991).

3. de Castro, L.F. et al. Activation of RANK/RANKL/OPG Pathway Is Involved in the Pathophysiology of Fibrous Dysplasia and Associated With Disease Burden. J Bone Miner Res 34, 290–294 (2019).

4. Zhao, X. et al. Expression of an active Galpha(s) mutant in skeletal stem cells is sufficient and necessary for fibrous dysplasia initiation and maintenance. Proc Natl Acad Sci U S A 115, E428–E437 (2018).

5. de Castro, L.F. et al. RANKL inhibition reduces lesional cellularity and Galpha(s) variant expression and enables osteogenic maturation in fibrous dysplasia. Bone Res 12, 10 (2024).

6. Szymczuk, V. et al. Safety and Efficacy of Moderate-dose Denosumab in Fibrous Dysplasia: Observational Results From a Phase 2 Clinical Trial. J Clin Endocrinol Metab 110, 2666–2673 (2025).

7. Liu, Z. et al. Protein kinase A is a dependent factor and therapeutic target in mouse models of fibrous dysplasia. Nat Commun 16, 5425 (2025).

8. Schalm, S.S. et al. Evaluation of Protein Kinase cAMP-Activated Catalytic Subunit Alpha as a Therapeutic Target for Fibrolamellar Carcinoma. Gastro Hep Adv 2, 307–321 (2023).

9. Bruystens, J.G.H. et al. A PKA-selective inhibitor captures an open but more ordered conformation of the PKA catalytic subunit. Proc Natl Acad Sci U S A 123, e2536312123 (2026).

10. Hanna, H., Mir, L.M. & Andre, F.M. In vitro osteoblastic differentiation of mesenchymal stem cells generates cell layers with distinct properties. Stem Cell Res Ther 9, 203 (2018).

11. Maridas, D.E., Rendina-Ruedy, E., Le, P.T. & Rosen, C.J. Isolation, Culture, and Differentiation of Bone Marrow Stromal Cells and Osteoclast Progenitors from Mice. J Vis Exp (2018).

12. Taylor, S.S., Ilouz, R., Zhang, P. & Kornev, A.P. Assembly of allosteric macromolecular switches: lessons from PKA. Nat Rev Mol Cell Biol 13, 646–658 (2012).

13. Kao, R.S. et al. Constitutive protein kinase A activity in osteocytes and late osteoblasts produces an anabolic effect on bone. Bone 55, 277–287 (2013).

14. Tascau, L. et al. Activation of Protein Kinase A in Mature Osteoblasts Promotes a Major Bone Anabolic Response. Endocrinology 157, 112–126 (2016).

15. Komori, T. Bone development by Hedgehog and Wnt signaling, Runx2, and Sp7. J Bone Miner Metab 43, 33–38 (2025).

16. Hwang, P.W. & Horton, J.A. Variable osteogenic performance of MC3T3-E1 subclones impacts their utility as models of osteoblast biology. Sci Rep 9, 8299 (2019).

17. Banno, Y., Sakai, T., Kumada, T. & Nozawa, Y. Potentiation by cholera toxin of bradykinin-induced inositol phosphate production in the osteoblast-like cell line MC3T3-E1. Biochem J 292 **(Pt** **2**), 401–408 (1993).

18. Katsianou, M.A. et al. The Critical Role of Transcription Factor RUNX2 in Bone Mechanobiology. Cells 15 (2025).

19. Thaweesapphithak, S. et al. Functional consequences of C-terminal mutations in RUNX2. Sci Rep 13, 12202 (2023).

20. Kim, H.J., Kim, W.J. & Ryoo, H.M. Post-Translational Regulations of Transcriptional Activity of RUNX2. Mol Cells 43, 160–167 (2020).

21. Sheridan, C.M., Heist, E.K., Beals, C.R., Crabtree, G.R. & Gardner, P. Protein kinase A negatively modulates the nuclear accumulation of NF-ATc1 by priming for subsequent phosphorylation by glycogen synthase kinase-3. J Biol Chem 277, 48664–48676 (2002).

22. Zhang, F., White, R.L. & Neufeld, K.L. Phosphorylation near nuclear localization signal regulates nuclear import of adenomatous polyposis coli protein. Proc Natl Acad Sci U S A 97, 12577–12582 (2000).

23. Brunet, A. et al. 14-3-3 transits to the nucleus and participates in dynamic nucleocytoplasmic transport. J Cell Biol 156, 817–828 (2002).

24. Shimizu, E., Nakatani, T., He, Z. & Partridge, N.C. Parathyroid hormone regulates histone deacetylase (HDAC) 4 through protein kinase A-mediated phosphorylation and dephosphorylation in osteoblastic cells. J Biol Chem 289, 21340–21350 (2014).

25. El-Masri, B.M. et al. Mapping RANKL-and OPG-expressing cells in bone tissue: the bone surface cells as activators of osteoclastogenesis and promoters of the denosumab rebound effect. Bone Res 12, 62 (2024).

26. Rashid, H., Smith, C.M., Convers, V., Clark, K. & Javed, A. Runx2 deletion in hypertrophic chondrocytes impairs osteoclast mediated bone resorption. Bone 181, 117014 (2024).

27. Wentworth, K.L. et al. Single-cell analysis of human fibrous dysplasia bone reveals a fibrotic transcriptome and GNAS variants in endothelial, perivascular, and stromal cells. Am J Hum Genet 112, 2067–2087 (2025).

28. Wang, D. et al. Isolation and characterization of MC3T3-E1 preosteoblast subclones with distinct in vitro and in vivo differentiation/mineralization potential. J Bone Miner Res 14, 893–903 (1999).

29. Charoenphandhu, N. et al. Responses of primary osteoblasts and osteoclasts from hemizygous beta-globin knockout thalassemic mice with elevated plasma glucose to 1,25-dihydroxyvitamin D(3). Sci Rep 9, 13963 (2019).

30. Subramanian, A. et al. Gene set enrichment analysis: a knowledge-based approach for interpreting genome-wide expression profiles. Proc Natl Acad Sci U S A 102, 15545–15550 (2005).

31. Zhou, Y. et al. Metascape provides a biologist-oriented resource for the analysis of systems-level datasets. Nat Commun 10, 1523 (2019).

32. Badia, I.M.P., et al. decoupleR: ensemble of computational methods to infer biological activities from omics data. Bioinform Adv 2, vbac016 (2022).

33. Muller-Dott, S. et al. Expanding the coverage of regulons from high-confidence prior knowledge for accurate estimation of transcription factor activities. Nucleic Acids Res 51, 10934–10949 (2023).

34. Teramoto, H. et al. The small GTP-binding protein rho activates c-Jun N-terminal kinases/stress-activated protein kinases in human kidney 293T cells. Evidence for a Pak-independent signaling pathway. J Biol Chem 271, 25731–25734 (1996).

35. Pierson Smela, M.D., et al. Directed differentiation of human iPSCs to functional ovarian granulosa-like cells via transcription factor overexpression. Elife 12 (2023).

36. Marinissen, M.J., Chiariello, M., Pallante, M. & Gutkind, J.S. A network of mitogen-activated protein kinases links G protein-coupled receptors to the c-jun promoter: a role for c-Jun NH2-terminal kinase, p38s, and extracellular signal-regulated kinase 5. Mol Cell Biol 19, 4289–4301 (1999).

37. Henriques-Oliveira, L. et al. Anchored screening identifies transcription factor blueprints underlying dendritic cell diversity and subset-specific anti-tumor immunity. Immunity 58, 2419–2438 e2413 (2025).

38. Ruiz-Saenz, A. et al. A reversible SRC-relayed COX2 inflammatory program drives resistance to BRAF and EGFR inhibition in BRAF(V600E) colorectal tumors. Nat Cancer 4, 240–256 (2023).

39. Ma, Y. et al. Making the invisible visible-ultrashort echo time magnetic resonance imaging: Technical developments and applications. Appl Phys Rev 9, 041303 (2022).

