## Supplementary figures for "A reversible GNAS-PKA-RUNX2 checkpoint governs skeletal stem cell fate in fibrous dysplasia"

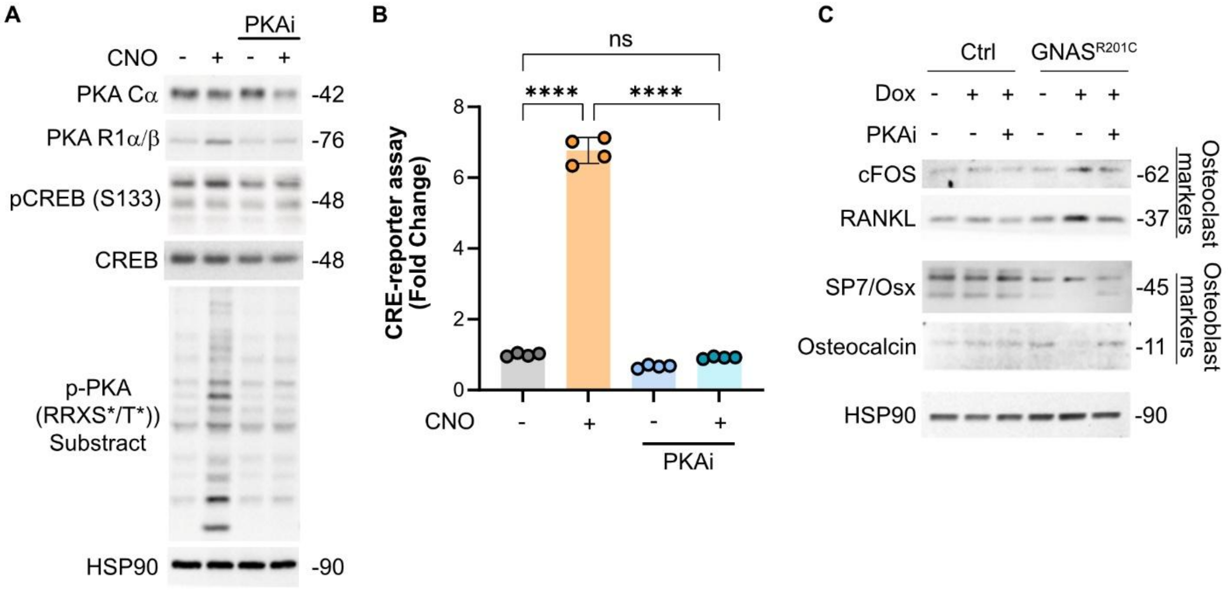


**Supplementary Figure S1. Pharmacological inhibition of PKA suppresses cAMP/PKA signaling and reverses GNAS$^{R201C}$-associated molecular changes. A.** Immunoblot analysis of cells treated with clozapine-N-oxide (CNO) in the presence or absence of a PKA inhibitor (PKAi). Protein levels of PKA Cα, PKA R1α/β, phosphorylated CREB (Ser133), total CREB, and phosphorylated PKA substrates (RRXS*/T*) were assessed. HSP90 served as a loading control. **B.** CRE-luciferase reporter activity in cells treated with CNO in the presence or absence of PKAi. Data are presented as fold change relative to untreated controls. **C.** Immunoblot analysis of control and GNAS$^{R201C}$ BMSCs treated with doxycycline (Dox) in the presence or absence of PKAi. Expression of osteoclast-associated markers (cFOS and RANKL) and osteoblast-associated markers (SP7/Osterix and Osteocalcin) was evaluated. HSP90 served as a loading control. Data are presented as mean ± SD. Each dot represents an independent biological replicate. One-way ANOVA determined statistical significance with multiple-comparison correction. ns, not significant; ****$P < 0.0001$.

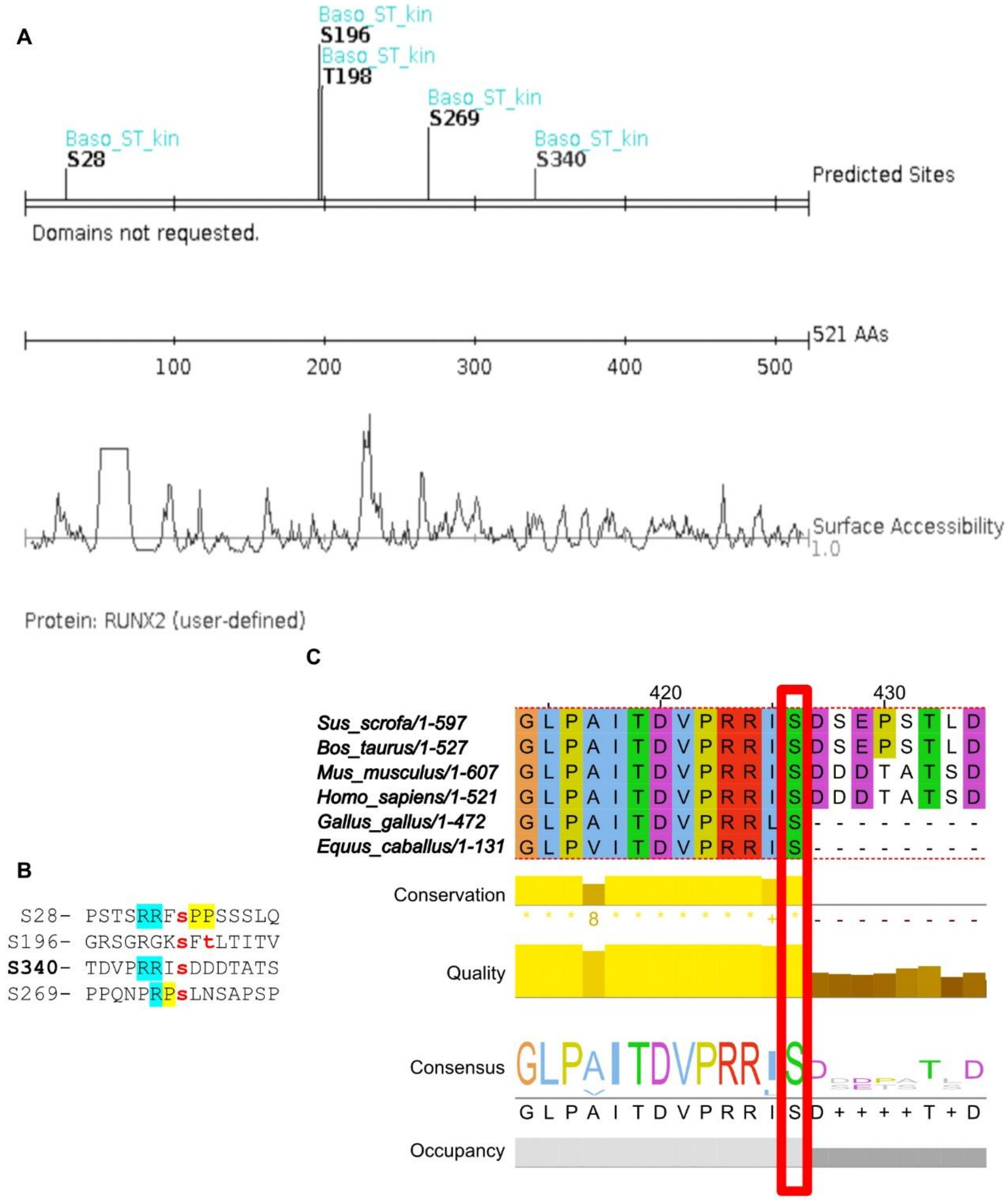


**Supplementary Figure S2. Identification of candidate PKA phosphorylation sites in RUNX2 and conservation of RUNX2 Ser340. A.** In silico prediction of serine/threonine kinase phosphorylation sites within RUNX2 by Scansite 4.0. Predicted PKA consensus motifs were identified at residues S28, S196, T198, S269, and S340. Surface accessibility analysis of the RUNX2 protein sequence is shown below. **B.** Amino acid sequences surrounding the predicted PKA phosphorylation sites in RUNX2. Basic residues characteristic of the PKA recognition motif are highlighted. **C.** Multiple sequence alignment of RUNX2 orthologs from *Sus scrofa*, *Bos taurus*, *Mus musculus*, *Homo sapiens*, *Gallus gallus*, and *Equus caballus*. The region encompassing Ser340 is highly conserved across species. The predicted PKA phosphorylation site at Ser340 is highlighted in red. Data were generated using bioinformatic analyses of the RUNX2 protein sequence.

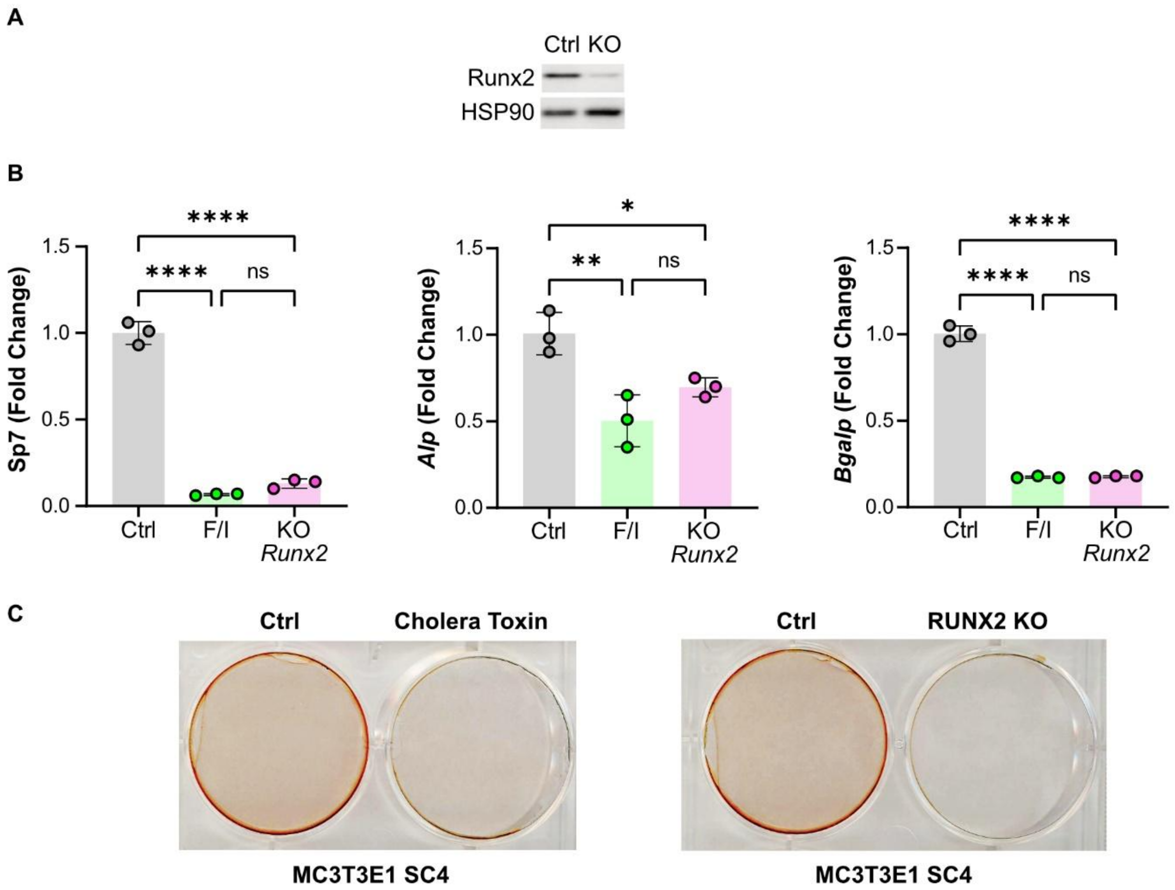


**Supplementary Figure S3. RUNX2 is required for osteoblast differentiation downstream of cAMP/PKA signaling. A.** Immunoblot validation of RUNX2 knockout (KO) MC3T3-E1 subclone 4 cells (MC cells). RUNX2 protein expression was assessed in control (Ctrl) and RUNX2 KO cells. HSP90 served as a loading control. **B.** Quantitative RT-PCR analysis of osteoblast-associated genes (*Sp7, Alpl,* and *Bglap*) in MC cells treated with forskolin/IBMX (F/I) or following RUNX2 knockout. Gene expression is presented relative to control cells. **C.** Representative Alizarin Red staining of MC cells following treatment with cholera toxin or genetic deletion of RUNX2. Mineralized matrix deposition was assessed after osteogenic differentiation. Data are presented as mean ±

SD. Each dot represents an independent biological replicate. One-way ANOVA determined statistical significance with multiple-comparison correction. ns, not significant; *P < 0.05, **P < 0.01, ****P < 0.0001.

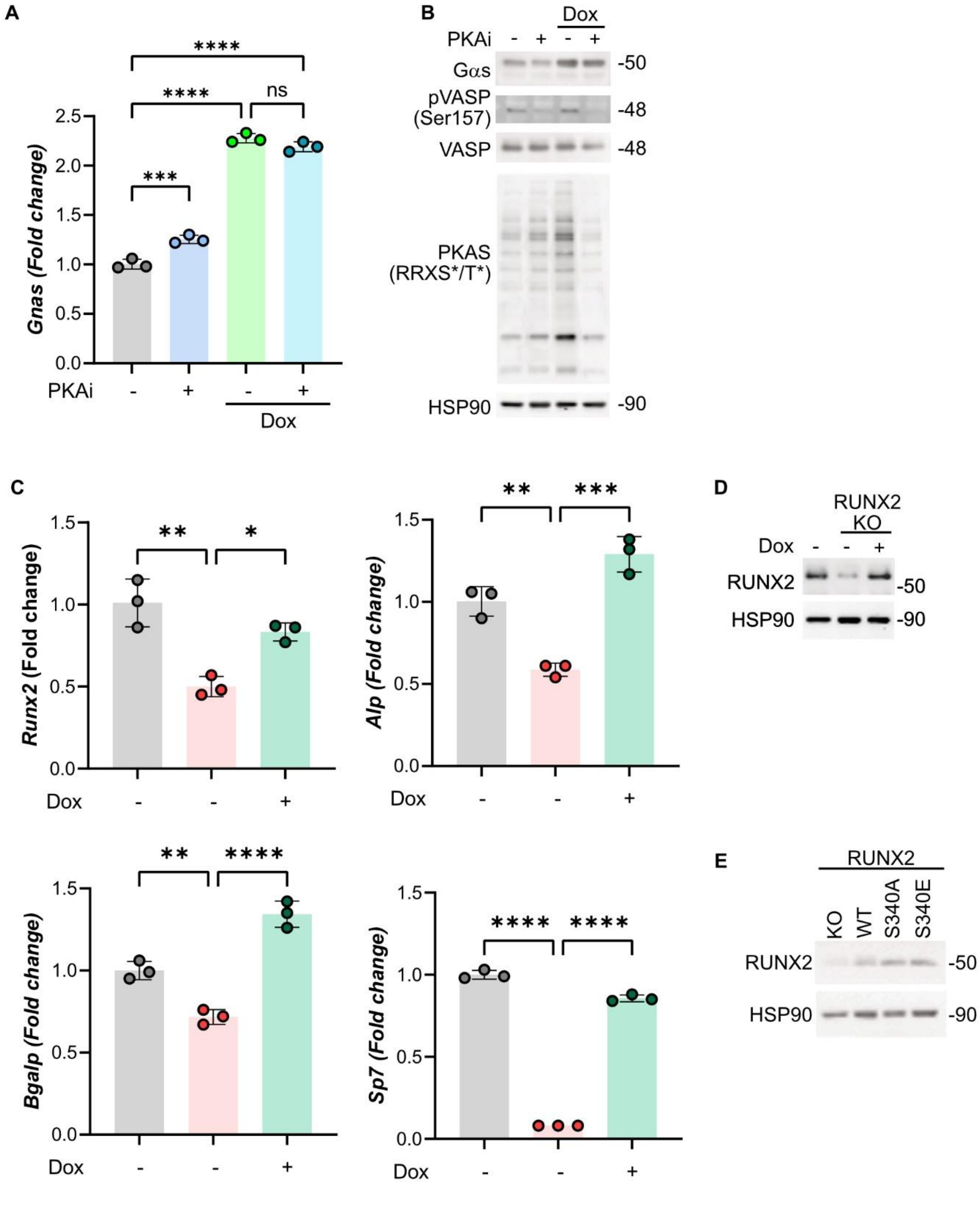

**Supplementary Figure S4. Restoration of RUNX2 expression rescues osteogenic gene expression in RUNX2-deficient cells: RUNX2 mutant analysis. A.** Quantitative RT-PCR analysis of *Gnas*$^{R201C}$ expression in FD BMSCs treated with doxycycline (Dox) in the presence or absence of a PKA inhibitor (PKAi). Dox-induced *Gnas*$^{R201C}$ expression was not significantly altered by PKA inhibition. **B.** Immunoblot analysis of FD BMSCs treated with Dox in the presence or absence of PKAi. Protein levels of Gαs, phosphorylated VASP (Ser157), total VASP, and phosphorylated PKA substrates (RRXS*/T*) were assessed. HSP90 served as a loading control. **C.** Quantitative RT-PCR analysis of osteoblast-associated genes (*Runx2, Alpl, Bglap,* and *Sp7*) in MC3T3-E1 subclone 4 RUNX2 knockout (KO) cells reconstituted with doxycycline (Dox)-inducible RUNX2-WT. Gene expression levels were compared between control, RUNX2 KO, and RUNX2-rescued cells. **D.** Immunoblot validation of doxycycline-inducible RUNX2-WT expression in MC3T3-E1 subclone 4 RUNX2 KO cells. HSP90 served as a loading control. **E.** Immunoblot validation of RUNX2 expression in MC3T3-E1 subclone 4 RUNX2 KO cells reconstituted with RUNX2-WT, RUNX2-S340A, or RUNX2-S340E expression constructs. HSP90 served as a loading control. Data are presented as mean ± SD. Each dot represents an independent biological replicate. One-way ANOVA determined statistical significance with multiple-comparison correction. ns, not significant; *$P$ < 0.05, **P < 0.01, ***P < 0.001, ****P < 0.0001.

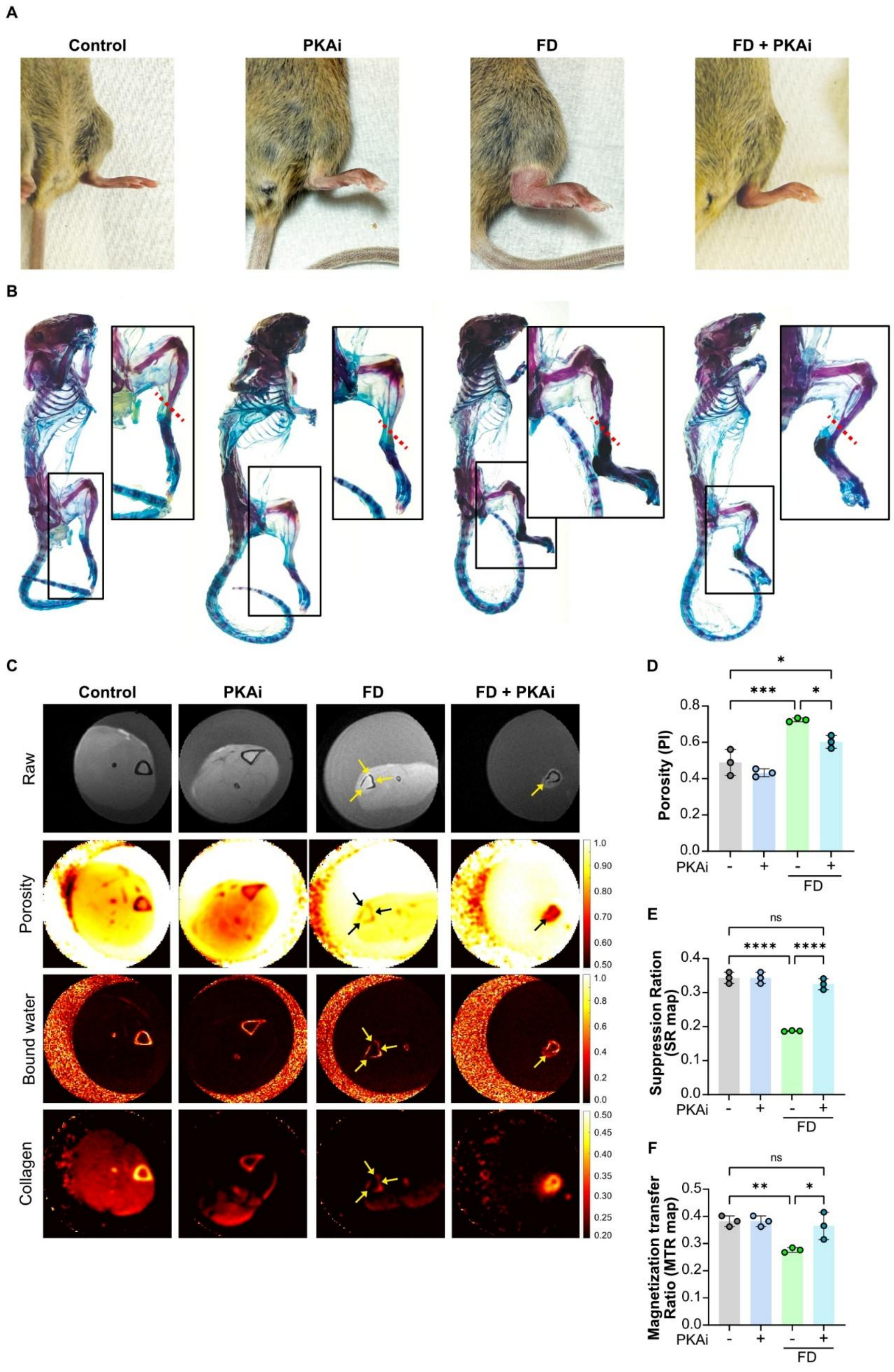
A
Control
PKAi
FD
FD + PKAi
B
C
Control
PKAi
FD
FD + PKAi
Raw
Porosity
Bound water
Collagen
D
Porosity (PI)
PKAi
FD
E
Suppression Ration (SR map)
ns
PKAi
FD
F
Magnetization transfer Ratio (MTR map)
ns
PKAi
FD

**Supplementary Figure S5. PKA inhibition decreases skeletal abnormalities and tissue defects in an FD mouse model. A.** Representative photographs of hindlimbs from control, PKA inhibitor (PKAi)-treated, FD, and FD+PKAi mice at the experimental endpoint. **B.** Representative whole-mount skeletal preparations from control, PKAi, FD, and FD+PKAi mice stained with Alcian Blue and Alizarin Red. Insets show higher magnification views of the affected hindlimb region. Red dashed lines indicate the anatomical location used for subsequent MRI analyses. **C.** Representative magnetic resonance imaging (MRI) analyses of hindlimb lesions from control, PKAi, FD, and FD+PKAi mice. Raw images, porosity maps, bound water maps, and collagen maps are shown. Arrows indicate fibrous dysplasia-associated lesions and regions of altered tissue architecture. **D.** Quantification of porosity index (PI) derived from MRI analyses in control mice. **E.** Quantification of suppression ratio obtained from solid-state MRI (SR map) analyses. **F.** Quantification of magnetization transfer ratio (MTR map) in control, PKAi, FD, and FD+PKAi mice. Data are presented as mean ± SD. Each dot represents an individual mouse. Statistical significance was determined by one-way ANOVA with multiple-comparison correction. ns, not significant; *P < 0.05, **P < 0.01, ***P < 0.001, ****P < 0.0001.
